# Single-cell resolution functional networks during unconsciousness are segregated into spatially intermixed modules

**DOI:** 10.1101/2023.09.14.557838

**Authors:** Daiki Kiyooka, Ikumi Oomoto, Jun Kitazono, Yoshihito Saito, Midori Kobayashi, Chie Matsubara, Kenta Kobayashi, Masanori Murayama, Masafumi Oizumi

## Abstract

The common neural mechanisms underlying the reduction of consciousness during sleep and anesthesia remain unclear. Previous studies have examined changes in network structure only using recordings with limited spatial resolution, which has hindered the investigation of the critical spatial scales from the micro (single neurons) to the meso (groups of neurons) for the reduction of consciousness. To address this issue, by leveraging fast, single-cell resolution, and wide-field two-photon microscopy, we recorded calcium signals from approximately 10,000 neurons across multiple cortical regions in awake, sleeping, and anesthetized mice. This single-cell resolution data enabled us to investigate the scales at which changes in network structure compared to an awake state are commonly observed during sleep and anesthesia. We found that at the single-cell scale, both sleep and anesthesia exhibited higher network modularity, indicating a segregated network, compared to an awake state. Despite this segregation, modules were spatially intermixed in all three states. In contrast, at the mesoscale, there were no consistent differences in modularity between states, and modules were spatially localized. Our multi-scale analysis provides novel insights into the cellular-scale organization of functional networks commonly associated with the reduction of consciousness and highlights a scale-dependent organization of network structures.

## Introduction

The process by which consciousness diminishes during sleep and anesthesia remains elusive in neuroscience. Prevailing theories of consciousness propose that the brain’s ability to integrate information is an indispensable prerequisite for consciousness, given the unified nature of the conscious experience [1–4]. Such information integration is enabled by the networked interactions of neurons at the microscale and brain regions at the macro/mesoscale. Therefore, it is crucial to examine to what extent and how the brain network is integrated or segregated at both the micro- and macro/mesoscale depending on the brain state [5–7]. Such network-level understanding is pivotal for elucidating the possibly shared neural underpinnings of reduced consciousness during sleep and anesthesia.

Current empirical findings from studies using macro/mesoscale neural recordings, such as functional MRI (fMRI) and electroencephalography (EEG), suggest that functional networks are more segregated during non-rapid eye movement (NREM) sleep and anesthesia than during an awake state [8–18]; that is, interactions across brain regions are diminished. For example, a perturbational approach using transcranial magnetic stimulation (TMS) has shown that TMS-evoked EEG potentials do not propagate across brain regions during NREM sleep, but do so during an awake state, implying a breakdown of cortical effective connectivity during NREM sleep [13]. Another approach, using a graph-theoretic measure called modularity, has shown that functional networks are more segregated into spatially localized modules during NREM sleep [10, 17] as well as during propofol-induced [19] and isoflurane-induced anesthesia [9, 16] than during an awake state.

While many past studies have investigated macro/mesoscale properties of networks [9, 10, 12, 16–18, 20–31], where the scales are mainly determined by the constraints of measurement technique, crucial spatial scales for understanding the mechanisms underlying the reduction of consciousness remain to be elucidated [4, 32–34]. To address this, it is essential to record neural activity at the single-cell resolution for two reasons. First, neurons represent one of the most fundamental units for information processing in the brain [35]. Second, by spatially coarse-graining single-cell resolution recordings, we can investigate macro/mesoscale properties of the network with signals directly derived from neural activity. The resulting macro/mesoscale data are qualitatively distinct from those obtained through conventional recording techniques, such as fMRI and EEG, which measure indirect proxies of neural activity. Although a few studies have investigated single-cell resolution functional networks, these studies have analyzed data from a limited brain area [36–40] or a limited number of neurons [41–43] because of the narrow spatial coverage of recording sites. These constraints make it difficult to investigate coarser spatial scales and cross-regional interactions, both of which are crucial for understanding brain network structures associated with the reduction of consciousness. Consequently, little is known about cellular-resolution functional networks across multi-modal brain regions and their relationship to networks at coarser spatial scales.

To uncover the changes in functional network structures commonly associated with the reduction of consciousness from the single-cell scale to the mesoscale, we recorded the activity of approximately 10,000 neurons during an awake state, sleep, and isoflurane-induced anesthesia using a fast, single-cell resolution, and wide-field-of-view (FOV) two-photon laser scanning microscope, FASHIO-2PM (fast-scanning high optical invariant two-photon microscopy), recently developed by our group [44, 45]. The FOV spanning a contiguous 3 mm *×* 3 mm area encompasses multiple cortical regions, including the somatosensory, motor, and other cortices, which enables us to explore the cellular-resolution functional networks over expansive brain areas. Importantly, by recording neuronal activity during sleep for the first time at the single-cell resolution across multi-modal brain regions, we can investigate changes in network structure shared between sleep and anesthesia compared to an awake state, revealing common structures associated with reduced consciousness.

To investigate to what extent and how the networks are segregated, we compare network modularity [46, 47] and the spatial distribution of modules during NREM sleep and anesthesia with those during an awake state, both at the single-cell and mesoscale levels. We show that modularity of single-cell resolution functional networks is higher during NREM sleep and anesthesia than during an awake state, with differences associated with mid-to-high-degree neurons rather than the highest-degree neurons. However, we show that these modules are distributed in a spatially intermixed manner in all states. On the other hand, modularity of mesoscale functional networks estimated from coarse-grained neural activity is not consistently different between states and the identified modules are spatially localized. Our results provide novel insights into the cellular-scale organization of functional networks universally associated with reduced consciousness and highlight a scale-dependent organization of network structures.

## Results

### Framework for investigating network structure

We investigate the structure of functional networks during an awake state, NREM sleep, and isoflurane-induced anesthesia particularly in terms of network segregation. Modularity is used as a measure to quantify the degree of network segregation. It quantifies the extent to which a network can be divided into non-overlapping “modules,” where neurons within the same modules are densely connected but those with different modules are sparsely connected. The lower the modularity, the more integrated the network, whereas the higher the modularity, the more segregated the network (Fig. 1a) [48] (see Materials and Methods for details).

**Figure 1:**
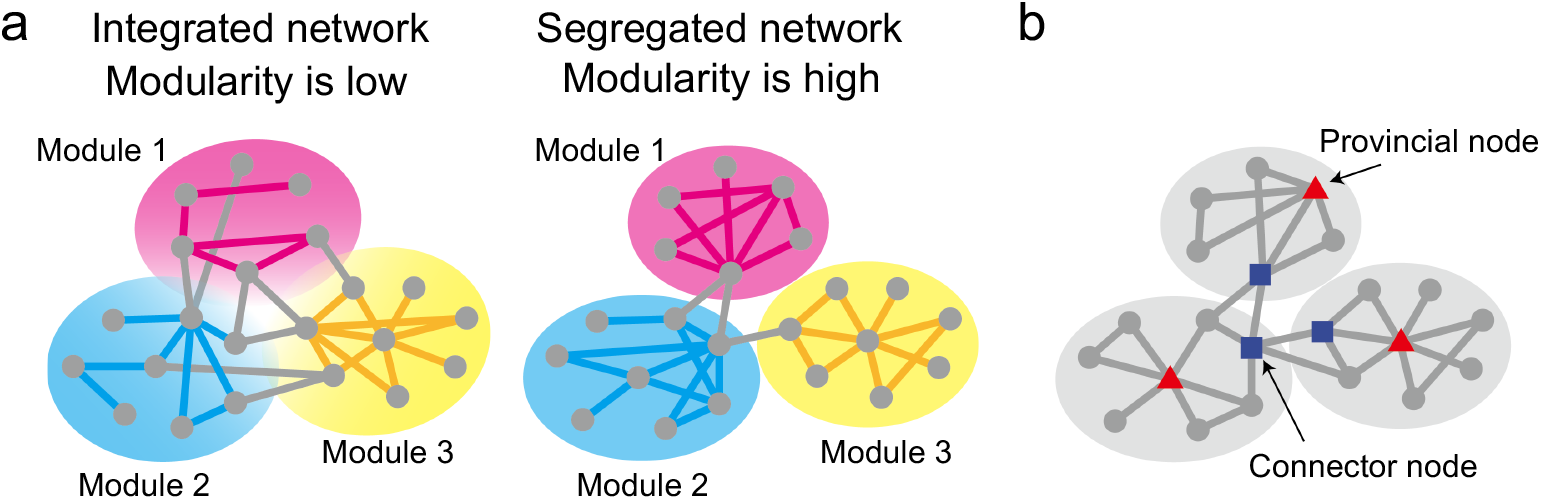
Schematic illustration of the concept of modularity (**a**) and two types of neurons (**b**).

Modularity is central to this study because it allows us to investigate not only the extent of segregation but also the manner in which networks are segregated or integrated. Calculating modularity involves identifying module assignments such that neurons within the same module have dense connections, while neurons in different modules have sparser connections. This approach gives us two key insights into how networks are segregated or integrated. First, we can characterize neurons based on the contribution to the modular structure of the networks. Neurons which have many across-module connections but few within-module connections are referred to as connector nodes and contribute to network integration, while neurons which have few across-module connections but many within-module connections are referred to as provincial nodes and contribute to network segregation (Fig. 1b). Analyzing these neurons provides a detailed single-cell perspective on how network segregation occurs. Second, we can examine the spatial distribution of the identified modules and gain insight into how functional networks are organized across brain regions.

### Fast, cell-resolution, contiguous-wide two-photon calcium imaging across multi-modal cortical areas during an awake state, sleep, and anesthesia

We recorded the cortical-wide activity of individual neurons in waking, sleeping, and anesthetized mice (Fig. 2), using FASHIO-2PM developed in our previous studies [44, 45]. The FOV spanning a contiguous 3 mm 3 *×* mm area allowed us to record multiple brain regions in cortical layer 2/3 excitatory neurons, including the somatosensory, motor and higher-order cortices (Fig. 2; see also Supplementary Video 1). To determine the sleep stages in mice, we simultaneously recorded electromyogram (EMG) potentials from neck muscles and EEG signals from screw electrodes placed on the lateral secondary visual area (V2L) of the imaged hemisphere (Fig. 2b) (see Materials and Methods). Four sleep states were determined by post-hoc analysis of the EEG and EMG: an awake state, NREM sleep, REM sleep, and a quiet awake state. As REM sleep and a quiet awake state were rare, we could not record long enough data to reliably estimate correlation during these states (see Table S1 for duration of these states). Therefore, we focused on an awake state and NREM sleep for subsequent analysis. To record neuronal activity during anesthesia, we anesthetized mice with 0.6% isoflurane inhalation after wake recording (hereafter, we used the terms sleep and anesthesia as NREM sleep and isoflurane-induced anesthesia, respectively, in the main text unless otherwise specified).

**Figure 2:**
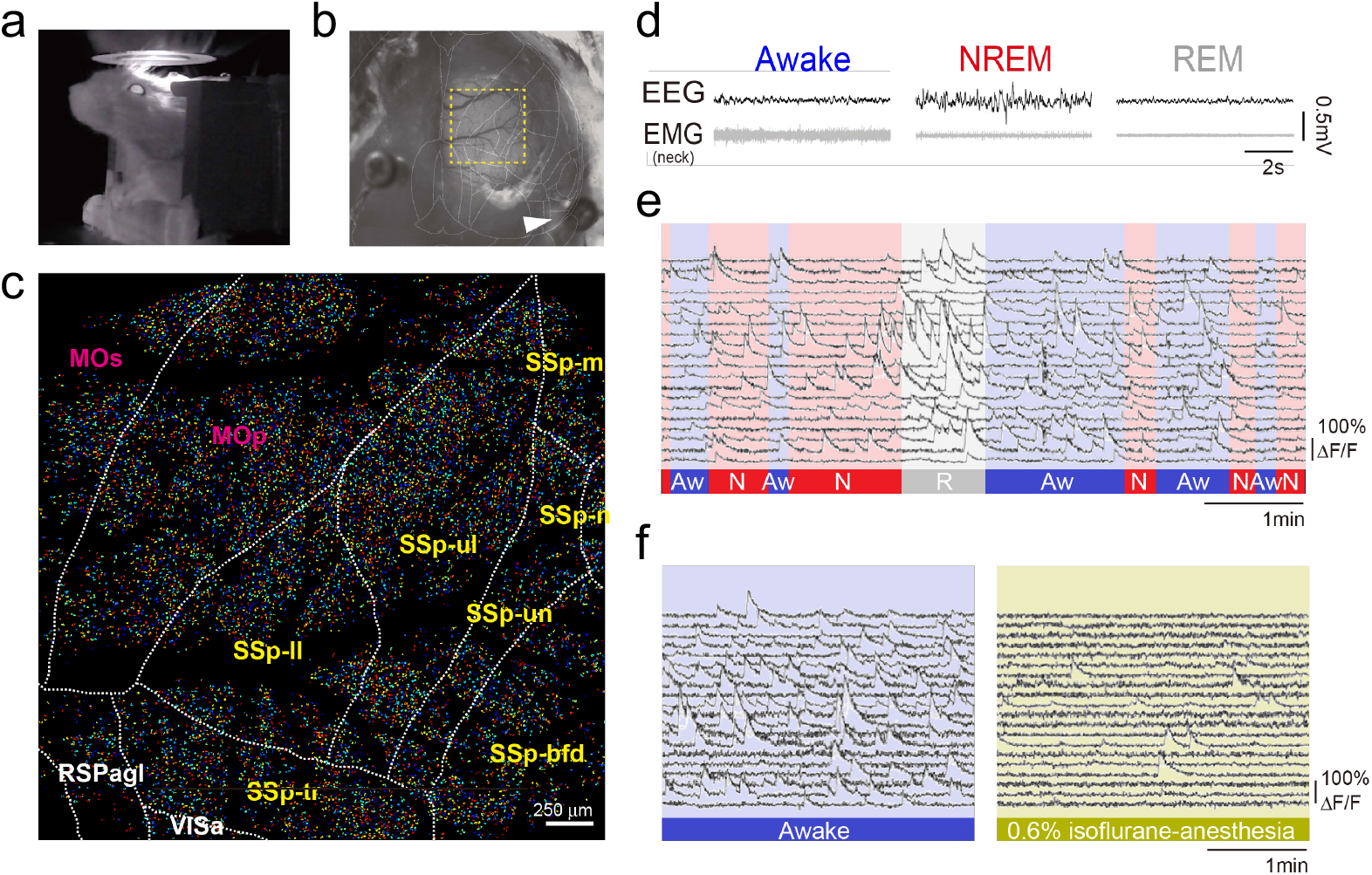
Fast, cell-resolution, wide-field two-photon calcium imaging from mice during an awake state and sleep. **a**. An infrared camera image of a mouse under the FASHIO-2PM. **b**. A dorsal view of the mouse head. A yellow dashed square indicates the field-of-view (FOV). An arrowhead indicates a screw to acquire EEG from the same hemisphere on the imaging side. Overlaid white lines show borders according to the Allen Mouse Common Coordinate Framework. **c**. Activity of *>* 10,000 neurons was recorded from a 3 mm 3*×* mm FOV that included multiple brain regions. Borders between the regions are shown as white dashed lines according to the Allen Mouse Common Coordinate Framework. MOp, primary motor; MOs, secondary motor; SSp-bfd, S1-barrel field; SSp-ll, S1-lower limb, SSp-m, S1-mouse; SSp-n, S1-nose; SSp-tr, S1-trunk; SSp-ul, S1-upper limb; SSp-un, S1-unassigned area; RSPagl, retrosplenial area, lateral agranular part. **d**. Examples of EEG and EMG recordings during the imaging session. The sleep-wake state was estimated by post-hoc analysis of EEG and EMG (see Materials and Methods). **e**. Representative Ca^2+^ signals from 20 randomly selected neurons are shown in sleep-wake states. The bottom color bar indicates the sleep-wake state of the mouse: Awake state (Aw, blue), NREM sleep (N, red), and REM sleep (R, gray). **f**. Representative Ca^2+^ signals from 20 randomly selected neurons are shown in awake or anesthetic states.

To prepare for network analysis, we investigated how the number of active neurons differed between brain states. This is because our network analysis focuses on neurons that have at least one Ca^2+^ event within all windows used to estimate the networks in each session, ensuring that a network size within a session is the same. We found that the number of neurons that have at least one Ca^2+^ event within each time-window was, on average, 65% during anesthesia, due to the reduction of neural activity caused by isoflurane inhalation, while it was above 95% during an awake state and sleep (Fig. S1; see also Figs. S2 and S3 for event frequency and mean amplitude of all mice). This led to the substantial reduction of the number of neurons used in the following network comparisons between an awake state and anesthesia (see Tables S1 and S2 for the number of neurons used for network analysis).

### Segregated functional networks during sleep and anesthesia

In this study, we separately compared functional networks during an awake state with those during sleep and with those during anesthesia. This is because wakefulness-sleep comparison and wakefulness- anesthesia comparison are based on recordings from the same session, whereas sleep and anesthesia were recorded in separate sessions, making direct comparisons between these states challenging. As mentioned earlier, because the number of neurons active during anesthesia is significantly smaller compared to an awake state and sleep, the number of neurons available for comparing network structures between an awake state and anesthesia is much smaller than that for an awake state and sleep. In the following, we investigate whether there are commonalities between sleep and anesthesia from the perspective of network modularity, despite significant differences in the level of neuronal activity between the two states and the number of neurons included in the analysis.

To compare modularity (1) during an awake state and sleep and (2) during an awake state and anesthesia, we estimated the functional networks in the three states. We calculated the Pearson correlation coefficients of the deconvolved Ca^2+^ signal between node pairs, i.e., neuron pairs (see Materials and Methods for details). We then binarized the correlation matrices at a threshold set such that the connection density was equal to a certain value, *K* (Fig. 3a, b) [49]. This procedure prevented network measures from being trivially different because of variations in the number of edges rather than in the pattern of connections (see Figs. S4 and S5 for the relationship between connection density and absolute correlation threshold during each state). To demonstrate the robustness of the results across a wide range of *K*, we varied *K* from 0.008 to 0.3, which we considered as the lowest and highest values, respectively, for valid network analysis. When *K* was 0.008, the threshold absolute correlation value in some mice exceeded 0.45, which could have led to the absence of edges in highly correlated neuron pairs. Thus, for *K* ≦ 0.008, we consider the networks unsuitable for graph-theoretic analysis. When *K* was 0.3, the threshold absolute correlation value in some mice was below 0.05, which can lead to the presence of edges between non-significantly correlated neuron pairs. Thus, for *K* ≧ 0.3, we consider the networks unsuitable for graph-theoretic analysis.

**Figure 3:**
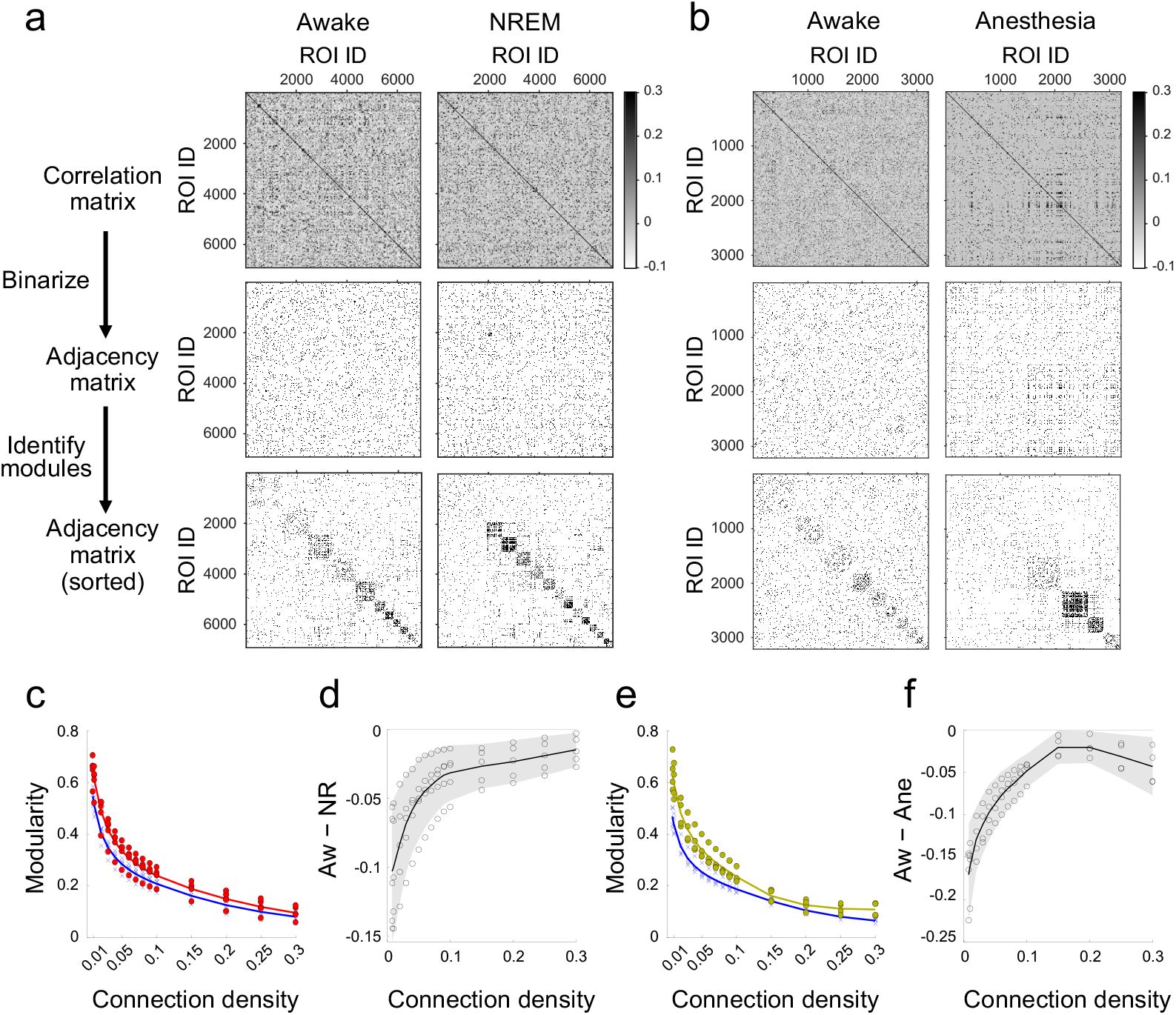
Estimation of functional networks and comparison of modularity. **a**. Procedures for estimating functional networks and calculating modularity during an awake state (left) and NREM sleep (right). **b**. Procedures for estimating functional networks and calculating modularity during an awake state (left) and anesthesia (right). **c**. Comparison of modularity between an awake state (blue) and NREM sleep (red). Crosses (awake) and dots (NREM sleep) represent modularity of individual mice, averaged across networks estimated from different time windows. Lines represent the mean modularity across mice. **d**. Difference in modularity between an awake state and NREM sleep. Each circle represents an individual mouse. The black line represents the average across mice. The gray-shaded area indicates the 95% confidence interval (based on one-sample t-test, *n* = 5, *df* = 4). **e**. Same as **c**, but comparing an awake state (blue) and anesthesia (yellow). **f**. Same as **d**, but comparing an awake state and anesthesia. The gray-shaded area indicates the 95% confidence interval (based on one-sample t-test, *n* = 4, *df* = 3).

To compute the modularity of the estimated functional networks, we first identified a module assignment that maximized the modularity index *Q* in Eq. (3), using the Louvain algorithm. The adjacency matrices, with neuron IDs sorted according to the identified modules, are shown in the bottom row of Fig. 3a, b. Each block along the diagonal corresponds to a module. The number of detected modules was not significantly different between an awake state and sleep across different connection densities (Fig. S6) (the 95% confidence intervals for the differences in the number of modules, shown as gray shaded areas in Fig. S6c, include zero across connection densities). The number of detected modules was not consistently different across mice between an awake state and anesthesia when 0.03 *≤ K*, but larger during anesthesia than during an awake state when *K <* 0.03 (Fig. S7). After module assignment, we computed the modularity using Eq. (3).

We found that modularity was higher during both sleep and anesthesia than during an awake state across different connection densities (Fig. 3c–f) (the 95% confidence intervals for the modularity differences, shown as gray shaded areas in Fig. 3d and f, remained negative across connection densities). This suggests that functional networks during sleep and anesthesia are more segregated into modules than those during an awake state. This difference could not be explained by the number of modules, because the number of modules was not significantly different between an awake state and sleep across all of the connection densities, nor between an awake state and anesthesia, except when *K <* 0.03 (Figs. S6 and S7). Furthermore, even when we set the number of modules to be the same (see Materials and Methods), modularity remained higher during sleep and anesthesia than during an awake state, although at *K* = 0.15 and 0.2 the variability of modularity differences between wakefulness and anesthesia became large, and the 95% confidence intervals included zero (Figs. S9 and S10). Therefore, sleep and anesthesia are qualitatively similar in terms of the extent of network segregation compared with an awake state.

To verify the robustness of the modularity results, we investigated other measures of network segregation. The most basic and widely used measures are global efficiency and transitivity. We found that global efficiency was higher during an awake state than during sleep and anesthesia, when *K <* 0.05, but not different when *K* ≧ 0.05 (Figs. S11 and S12). We also found that transitivity was higher during sleep and anesthesia than during an awake state across different connection densities (Figs. S13 and S14). These results also suggest that functional networks are more segregated during both sleep and anesthesia than during an awake state. This indicates the robustness of the modularity results.

### Highest-degree neurons do not necessarily contribute to differences in modularity

Although we found in the previous section that sleep and anesthesia exhibited similar overall network trends in terms of increased modularity, it remains unclear whether individual neurons contribute to modularity differences in a similar manner. Specifically, the observed increase in overall modularity may result from an increase in the number of provincial nodes, a decrease in the number of connector nodes, or a combination of both. Furthermore, it is also unclear whether these differences result from the influence of a small number of highest-degree neurons or from the broader population of neurons.

To investigate each neuron’s contribution to modularity, we propose a node-specific modularity measure, *Q*_*i*_, by decomposing the overall modularity, *Q*, into contributions of each node:

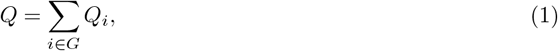

where *G* is the set of all nodes in the network. This measure quantifies the extent to which a node *i* contributes to the modularity *Q* (See Materials and Methods for details) and is defined as

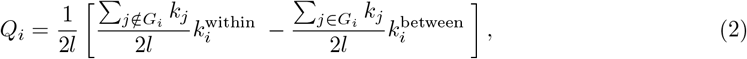

where *l* is the number of edges in the network, *G*_*i*_ is the set of nodes that belong to the same module as node *i, k*_*j*_ is the degree of node *j*,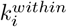 is the within-module degree of a node *i*, and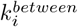is the across-module degree of a node *i*. Intuitively, the higher the within-module degree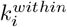 and the lower the across-module degree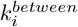, the higher the *Q*_*i*_, which indicates a greater contribution to modularity. In other words, neurons with high *Q*_*i*_ play a provincial-node-like role, whereas neurons with low *Q*_*i*_ play a connector-node-like role (Fig. 1b).

To determine whether the increases in modularity during sleep and anesthesia are associated with changes in the number of provincial or connector nodes, we compared the histogram of *Q*_*i*_ between an awake state and sleep as well as between an awake state and anesthesia. We found that the number of neurons that exhibited high *Q*_*i*_ values (neurons with *Q*_*i*_ larger than approximately 0.5 *×* 10^*−*4^ in Fig. 4a) was larger during sleep than during an awake state, while the number of neurons that exhibited low *Q*_*i*_ values (neurons with *Q*_*i*_ smaller than approximately 0.5 × 10^*−*4^ in Fig. 4a) was smaller during sleep than during an awake state (Fig. 4a for representative data; Fig. S15a–c for other connection densities and other mice). Similarly, the number of neurons that exhibited high *Q*_*i*_ values (neurons with *Q*_*i*_ larger than approximately 1.0 *×* 10^*−*4^ in Fig. 4b) was larger during anesthesia than during an awake state, while the number of neurons that exhibited low *Q*_*i*_ values (neurons with *Q*_*i*_ smaller than approximately 1.0*×*10^*−*4^ in Fig. 4b) was smaller during anesthesia than during an awake state (Fig. 4b for representative data; Fig. S15d–f for other connection densities and other mice). These results suggest that functional networks during both sleep and anesthesia exhibit similar segregation patterns, characterized by a decrease in the number of connector neurons and an increase in the number of provincial neurons.

**Figure 4:**
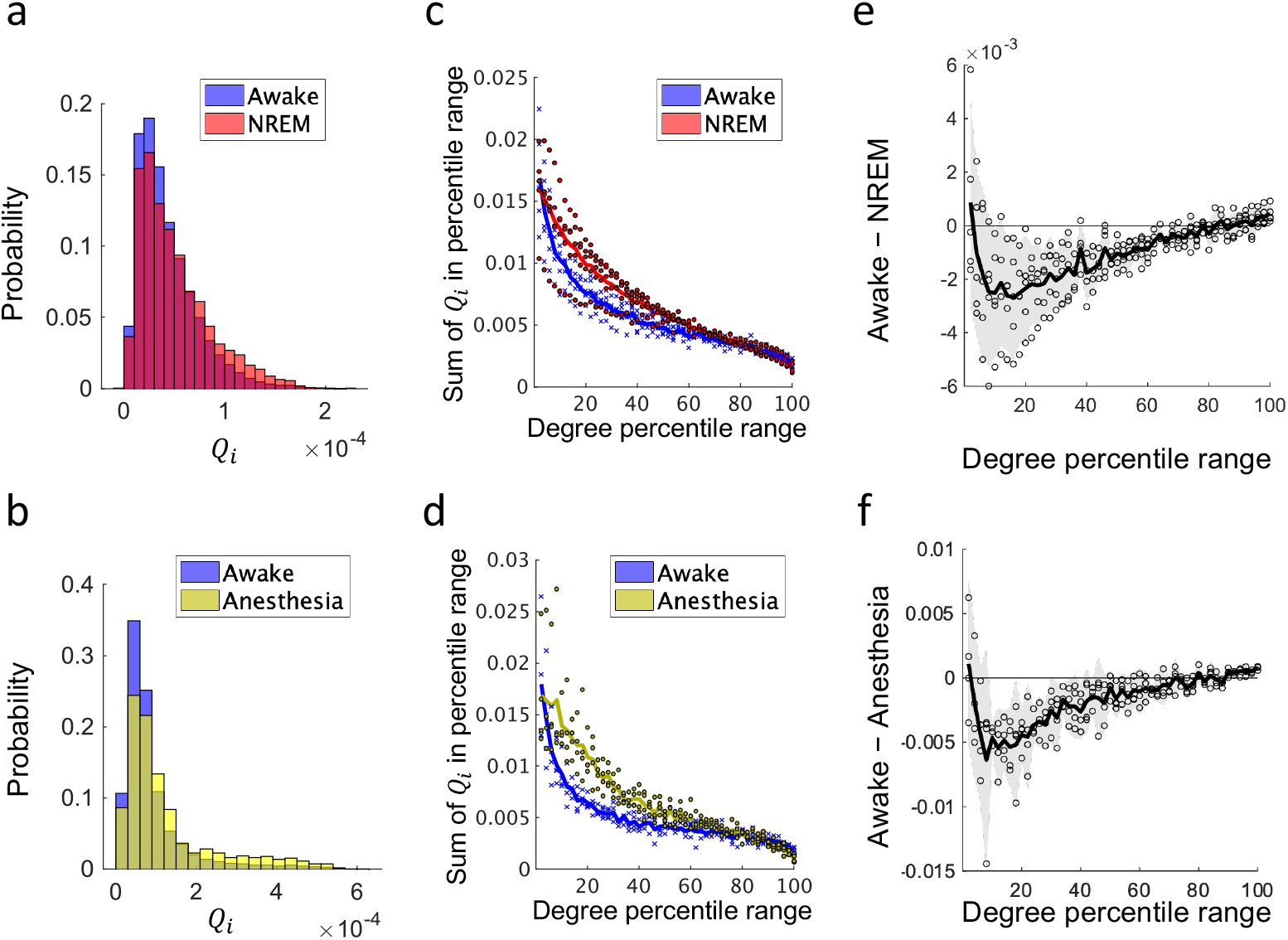
Contribution of each neuron to modularity (*K* = 0.05). For the data of different *K*, see Figs. S15 and S16. **a**. Representative histogram of *Q*_*i*_ during an awake state (blue) and NREM sleep (red). See also Fig. S15 for the histograms of all mice. **b**. Representative histogram of *Q*_*i*_ during an awake state (blue) and anesthesia (yellow). See also Fig. S15 for the histograms of all mice. **c**. Comparison of contributions to modularity across degree percentiles between an awake state (blue) and NREM sleep (red). For a given percentile range (x-axis value *x*), the sum of *Q*_*i*_ across neurons ranked in the top *x−*2% to *x*% by degree are plotted. Crosses (an awake state) and dots (NREM sleep) represent these percentile-based sums of *Q*_*i*_ for individual mice, averaged across networks estimated from different time windows. Lines represent the mean value across mice. **d**. Same as **c**, but comparing an awake state (blue) and anesthesia (yellow). **e**. Difference in the percentile-based sum of modularity between an awake state and NREM sleep. Each circle represents an individual mouse. The black line represents the average across mice. The gray-shaded area indicates the 95% confidence interval (based on one-sample t-test, *n* = 5, *df* = 4). **f**. Same as **e**, but comparing an awake state and anesthesia. The gray-shaded area indicates the 95% confidence interval (based on one-sample t-test, *n* = 4, *df* = 3).

To further investigate individual contributions to state-dependent modularity differences, we examined the relationship between *Q*_*i*_ and the degree. Contrary to the conventional focus on hub neurons (those with the highest degrees), we found that neurons with degrees in the top 10–60% percentile, rather than the top 10%, were primarily responsible for the differences in modularity between an awake state and sleep when *K* = 0.05 (Fig. 4c, e) (Fig. 4e shows that the 95% confidence interval for the difference in the contribution to modularity, based on one-sample t-test, *n* = 5, *df* = 4 is negative within the 10–60% percentile range). Likewise, neurons with degrees in the top 10–40% percentile, rather than the top 10% percentile, were primarily responsible for the differences in modularity between an awake state and anesthesia when *K* = 0.05 (Fig. 4d, f) (Fig. 4f shows that the 95% confidence interval for the difference in the contribution to modularity, based on one-sample t-test, *n* = 4, *df* = 3 is negative within the 10–60% percentile range). Although the specific percentile ranges that contribute to modularity differences varied depending on *K*, the highest-degree neurons were not consistently related to modularity differences across different *K* values (Fig. S16). These results suggest that sleep and anesthesia exhibit a similar pattern in which mid-to-high-degree neurons, rather than the highest-degree neurons, are primarily associated with the observed differences in modularity relative to an awake state.

### Spatially intermixed modules of single-cell resolution functional networks

To examine whether modules are spatially intermixed or localized on the cortex, we examined the spatial distribution of modules. We found that the modules were spatially intermixed during an awake state (Fig. 5a), sleep (Fig. 5b), and anesthesia (Fig. 5c). This indicates that adjacent neurons do not necessarily belong to the same module, while distant neurons sometimes belong to the same module (left column in Fig. 5a–c), although there is a little spatial imbalance of the distribution of modules. Thus, even during sleep and anesthesia, when functional networks are “segregated”, long-range functional connections remain across brain regions. This spatial distribution was consistent across all states, although the degree of segregation differed between an awake state and sleep as well as between an awake state and anesthesia.

**Figure 5:**
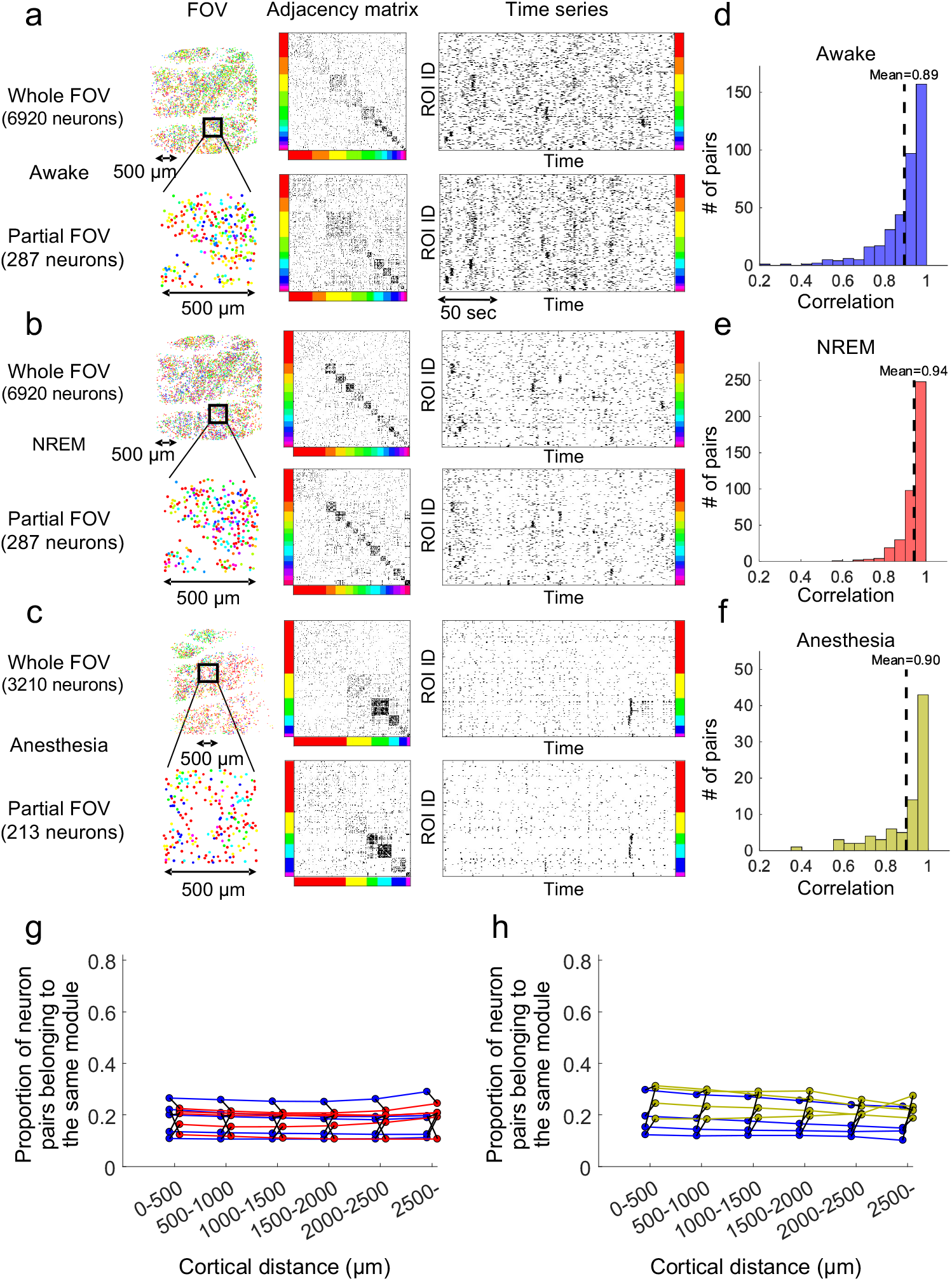
Spatially intermixed distribution of single-cell resolution functional network modules (*K* = 0.05). **a**. Spatial distribution of modules (left), adjacency matrix (middle), and time series (right) for both the entire FOV (top) and the partial FOV (bottom) of representative functional networks during an awake state. Figure 5: Neurons within the adjacency matrices and time series are sorted by their respective modules. Each color represents a specific module. **b**. Same as **a**, but during NREM sleep. **c**. Same as **a**, but during anesthesia. **d–f**. Histogram of correlation of module composition between the entire FOV and partial FOVs during an awake state (**d**), NREM sleep (**e**), and anesthesia (**f**). Each histogram shows data of all windows across all mice. Dotted lines represent the mean correlation. For the data of other *K*, see Fig. S17a, c. **g, h**. Relationships between cortical distance and module assignments of single-cell resolution functional networks (*K* = 0.05) during an awake state (blue), NREM sleep (red), and anesthesia (yellow). The x-axis represents the cortical distance and the y-axis represents the proportion of neuron pairs belonging to the same modules. Each dot represents an individual mouse, with data averaged across networks estimated from different time windows. Data from the same mouse is connected with lines. For the data of other *K*, see Fig. S17b, d.

We next quantitatively confirmed the above result in two ways. First, we calculated the correlation of the module proportion between the partial FOVs and the entire FOV. For the awake state/sleep data, we used non-overlapping 0.5 mm*×* 0.5 mm regions containing more than 250 neurons as partial FOVs. For the awake state/anesthesia data, due to the smaller number of analyzed neurons, we used non-overlapping 0.5 mm *×* 0.5 mm regions containing more than 100 neurons as partial FOVs. The correlation was on average higher than 0.89 during all states (*K* = 0.05) (Fig. 5d–f; Fig. S17a, c for different *K*). This indicates that the module composition within the most partial FOVs is similar to that of the entire FOV. Second, we examined the relationships between cortical distance and the probability of two neurons belonging to the same module. We found that the probability did not depend on cortical distance (Fig. 5g, h for *K* = 0.05, Fig. S17b, d for different *K*). This suggests that nearby neurons do not necessarily belong to the same module, while distant neurons sometimes do.

### Stability of modules

To investigate whether spatially intermixed modules identified in the previous section are randomly distributed or possess a temporally stable structure, we calculated the similarity of module assignments between different time windows. Initially, we computed the proportion of neuron pairs that were consistently assigned to the same module across two windows. We found that matching rates were on average 60–80 % (*K* = 0.05) (Fig. 6a, b; Fig. S18a, b, e, f for other *K*). To evaluate the significance of these matching rates, we generated a null distribution by randomly shuffling module assignments and calculating the matching rates 100 times. We then computed a z-score by dividing the difference between the observed matching rate and the mean matching rate of the null distribution by the standard deviation of the null distribution’s matching rates. We found that for both an awake state and sleep, the z-scores of module similarity were consistently greater than 3 (gray dashed line in Fig. 6c) across all mice, regardless of whether the comparison was within or between states (Fig. 6c; Fig. S18c, g for other *K*). Similarly, for an awake state and anesthesia, within-state comparisons yielded z-scores greater than 3 for all mice, and across-state comparisons exceeded this threshold in three out of four mice (Fig. 6d; Fig. S18d, h for other *K*). These findings demonstrate that, despite the spatially intermixed distribution of modules, these neural ensembles exhibit a temporal stability. In other words, they are not randomly distributed but possess a structured organization.

**Figure 6:**
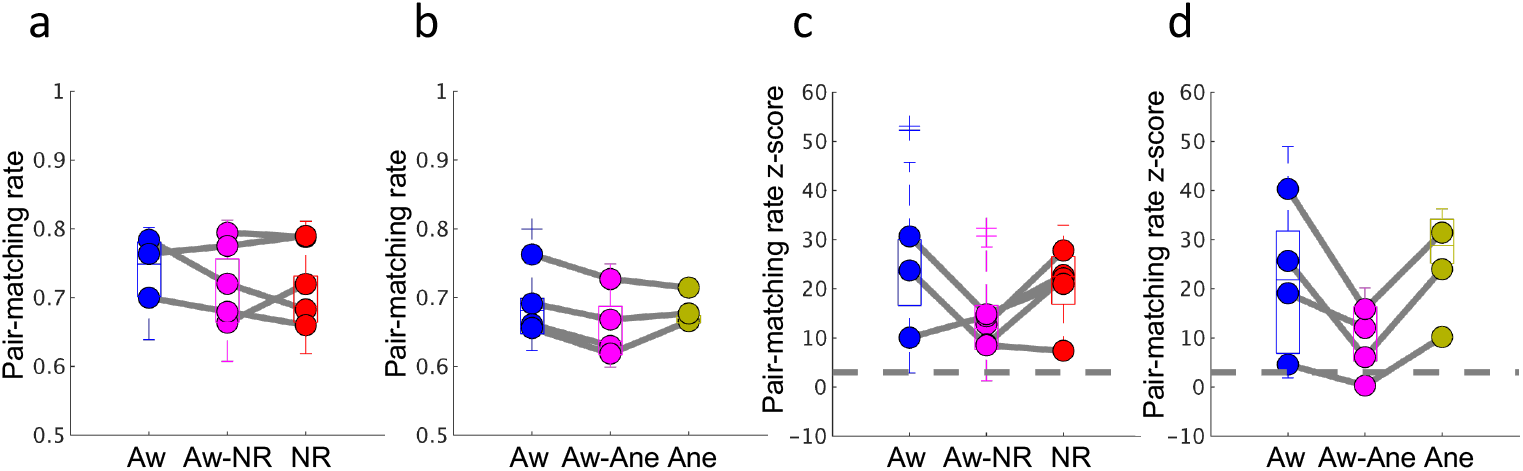
Stability of modules. **a, b**. The pair-matching rate represents the proportion of neuron pairs consistently assigned to the same module across two windows. “Aw”, “NR”, and “Ane” represent the “Aw-Ane” represent the stability of modules between an awake state and NREM sleep and between an awake state and anesthesia, respectively. Each dot corresponds to an individual mouse, with the value averaged across networks estimated from different time windows. In within-state comparisons, data without dots correspond to individuals where only one time window was available. The boxplots show data of all windows without distinguishing individuals, and crosses represent outliers. **c, d**. The z-score of **a** and **b**. The gray dashed lines represent a z-score value of 3.

### Modularity and distribution of modules at different spatial scales

To investigate how modularity and the spatial distribution of modules change depending on spatial scales, we spatially coarse-grained cellular-scale deconvolved Ca^2+^ signals and estimated functional networks (Fig. 7a). For the spatial coarse-graining, we clustered every *n*_*nei*_ neighboring neurons into a single parcel, where *n*_*nei*_ ranged from 1 (without coarse-graining) to 160. We then averaged the deconvolved Ca^2+^ signals within each parcel, and computed the Pearson correlation between parcels. Subsequently, we binarized the resulting correlation matrices and calculated modularity.

**Figure 7:**
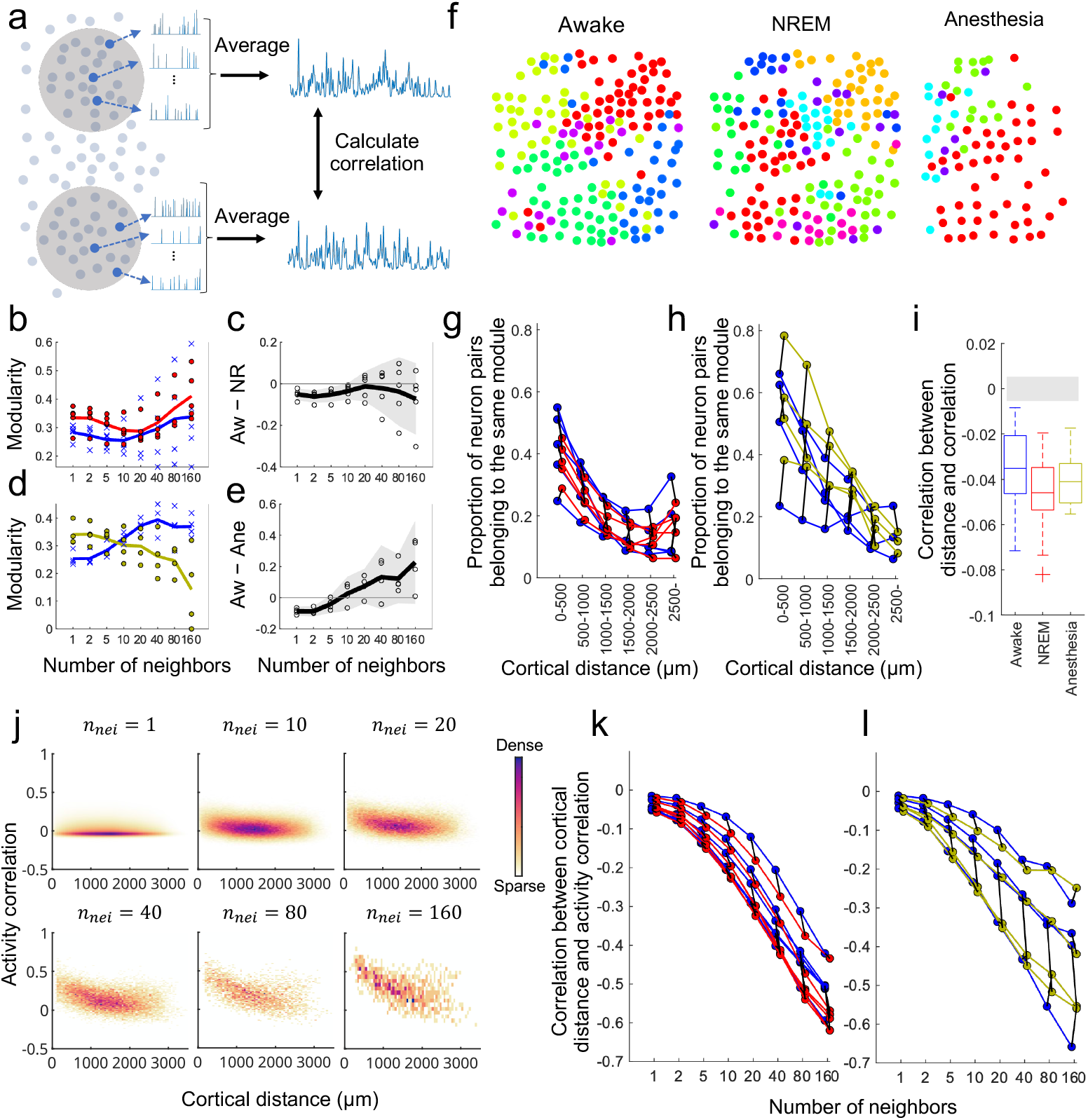
Modularity and distribution of modules at different spatial scales. **a**. Schematics of coarsegrained mesoscale functional network estimation. **b**. Comparison of modularity of functional networks between an awake state (blue) and NREM sleep (red) at different spatial scales (*K* = 0.05). Crosses (an awake state) and dots (NREM sleep) represent modularity of individual mice, averaged across networks estimated from different time windows. Lines represent the mean modularity across mice. **c**. Difference in modularity between an awake state and NREM sleep at different spatial scales (*K* = 0.05). Each circle represents an individual mouse. The black line represents the average across mice. The gray-shaded area indicates the 95% confidence interval (based on one-sample t-test, *n* = 5, *df* = 4). **d**. Same as **b**, but comparing an awake state (blue) and anesthesia (yellow). **e**. Same as **c**, but comparing an awake state and anesthesia. The gray-shaded area indicates the 95% confidence interval (based on one-sample t-test, *n* = 4, *df* = 3). Figure 7: **f**. Example of the spatial distribution of coarse-grained functional network modules during each state (*K* = 0.05, *n*_*nei*_ = 40). Each color represents a specific module. **g, h**. Relationships between cortical distance and module assignments of coarse-grained functional networks (*K* = 0.05, *n*_*nei*_ = 40) during an awake state (blue), NREM sleep (red), and anesthesia (yellow). The coordinates of each parcel used for distance calculation are defined as the average coordinates of the neurons that constitute the parcel. Each dot represents an individual mouse, with data averaged across networks estimated from different time windows. Data from the same mouse is connected with lines. **i**. Correlation between cortical distance and activity correlation during each state. The boxplots include all windows across all mice. The gray-shaded area represents the range between the upper and lower 2.5 percentiles of the correlation between distance and activity correlation when the position information is shuffled. This percentile range is calculated for each window by generating 1000 null model correlations, and the strictest range across all windows is illustrated. **j**. Example of the relationship between cortical distance and activity correlation (during an awake state) at different levels of coarse-graining. **k, l**. Correlation between cortical distance and activity correlation during an awake state (blue), NREM sleep (red), and anesthesia (yellow) at different levels of coarse-graining. Each dot represents an individual mouse, with data averaged across networks estimated from different time windows. Data from the same mouse is connected with lines.

We found that as the level of coarse-graining increased, network modularity did not show consistent differences across different mice between an awake state and sleep (Fig. 7b, c) nor between an awake state and anesthesia (Fig. 7d, e) (see also Fig. S19 for changes in modularity values at each spatial scale as a function of *K*). Specifically, for the comparison between an awake state and sleep, the variability of modularity differences became large, and the 95% confidence intervals included zero when 10 *≤ n*_*nei*_ (gray-shaded area in Fig. 7c). In the comparison between an awake state and anesthesia, the modularity values were, on average, higher during an awake state than anesthesia when *n*_*nei*_ was large (Fig. 7d, e). However, the variability was substantial across mice, and the 95% confidence intervals included zero when 5 *≤ n*_*nei*_(gray-shaded area in Fig. 7e).

As for the spatial distribution of the modules, we found that the modules were localized during all states in the coarse-grained mesoscale networks (Fig. 7f). In other words, unlike cellular-resolution networks (Fig. 5), nearby parcels were more likely to belong to the same module than distant ones stability of modules within an awake state, NREM sleep, and anesthesia, respectively. “Aw-NR” and

(Fig. 7g, h for *K* = 0.05, *n*_*nei*_ = 40; Fig. S20 for other *K* values).

One possible explanation for the change in module distribution with coarse-graining is that correlations between nearby neurons were slightly stronger than those between distant neurons but not strong enough for the module distribution to be localized at the cellular scale. If this were the case, as correlations build up through coarse graining, the correlations between nearby parcels would become markedly higher than those between distant parcels, leading to module localization.

We investigated how the correlation depended on the distance at each spatial scale and confirmed that the above explanation was indeed the case. First, we calculated correlation between cortical distance and activity correlation at the single-cell level and compared it with that of shuffled data. We found a subtle but significant dependence of the correlation on distance during all states (all data points in Fig. 7i are below the gray-shaded area) even at the single-cell level. Second, we investigated how the distance dependency of activity correlation changed when we coarse-grained neural activity into different spatial scales. We found that as the level of coarse-graining increased, the distance dependency of activity correlation became more intense during all states due to the buildup of weak correlations at the cellular level (Fig. 7j–l). This intensified dependence accounts for the module localization in coarse-grained networks.

## Discussion

In this study, we recorded the spontaneous activities of approximately 10,000 neurons in a contiguous 3 mm 3 *×*mm FOV, including multiple cortices in each mouse (Fig. 2) and compared the structure of functional networks between an awake state and sleep, as well as between an awake state and anesthesia, at both the single-cell and mesoscale levels. At the single-cell level, we found that modularity was higher during both sleep and anesthesia than an awake state (Fig. 3c–f), and that the contribution of the highest-degree neurons to modularity did not necessarily differ between an awake state and sleep, or an awake state and anesthesia, while mid-to-high-degree neurons contributed more to the modularity difference (Fig. 4c–f). We also found that modules were spatially intermixed during all states (Fig. 5), yet exhibited a temporal stability (Fig. 6). Unlike cellular-resolution functional networks, mesoscale functional networks estimated from coarse-grained neural activity did not show statistically robust differences in modularity between states (7b–e) and modules were spatially localized during all states (Fig. 7f–h).

### Segregation of functional networks during sleep and anesthesia at the cellular scale

While previous studies using macro/mesoscale recordings have already shown that network segregation increases during both sleep and anesthesia [9, 10, 16–19], our findings on the cellular-resolution functional networks represent a conceptually novel contribution rather than an incremental result. This is because the structure of single-cell resolution functional networks cannot be inferred from macro/mesoscale observations alone, while the reverse is possible, i.e., the properties of macro/mesoscale functional networks can be inferred by coarse-graining the single-cell resolution data. Therefore, network segregation observed at the single-cell level provides fundamentally new insights that cannot be extrapolated from macroscale functional networks.

In addition, our result on the decomposition of modularity offers an alternative perspective to brain network studies that focus on hub nodes, those with the highest degrees [50–52]. In general, the highest-degree hubs are thought to play significant roles in the network. Given that an awake and sleep/anesthesia are physiologically and functionally different, one might expect that the highest-degree neurons would have a large effect on the change in modularity depending on the level of consciousness. However, we found that the differences in modularity between an awake state and sleep as well as between an awake state and anesthesia were largely associated with mid-to-high-degree rather than the highest-degree neurons. These results suggest that the broader population of neurons contributes to network segregation. It should be noted that this does not mean that the highest-degree neurons play no part in the brain function. Rather, these neurons may serve fundamental roles necessary for maintaining essential brain functions regardless of consciousness state, which could explain why their contribution to modularity remains relatively stable. Our results highlight the importance of investigating network structure not only through highest-degree hub neurons but also by considering contributions across all neurons. Therefore, our findings suggest that a more inclusive approach to network structure is essential for understanding brain function at a single-cell level. Such an approach is not only relevant to consciousness research but also holds promise for exploring pathological brain conditions from a novel perspective. For instance, while brain networks in individuals with Alzheimer’s disease (AD) are known to exhibit disturbed modular structures [53, 54] and changes in hub regions [55–58], future studies could examine how degree-specific contributions to modularity evolve with AD progression. Such investigations could provide a more comprehensive understanding of brain diseases and their relationships to network structures.

### Neural mechanisms underlying spatially intermixed modules

Our finding of spatially intermixed modules brings about conceptual advancement by challenging the conventional view on how brain regions interact during unconsciousness [13]. As for spatially intermixing correlation patterns, previous studies utilizing single-cell resolution recording techniques have reported that neurons with different stimulus selectivity are distributed in a spatially intermixed manner within a single modality such as the visual [59] and sensory cortex [60]. The novelty of our finding is that modules are intermixed not only within but also across multi-modal brain regions. Since spatially intermixed modules imply that distant neurons sometimes belong to the same module, our finding suggests that not only during an awake state but also during sleep and anesthesia, when functional networks are “segregated”, long-range functional connections remain across multi-modal brain regions. This is in contrast to conventionally studied macro/mesoscale functional networks, where modules exhibit spatial localization [17, 23, 30, 61]. Such nuanced segregation can only be discovered through simultaneously recording the neural activity during different brain states across multi-modal brain regions at the single-cell resolution.

There are at least three possible mechanisms underlying spatially intermixed distribution of modules: common inputs, poly-synaptic inputs, and strong and variable recurrent interactions. First, if there are common inputs to distant neurons and such common inputs do not depend much on the distance between the neurons, we would observe spatially intermixed modules. Consistent with this, recent empirical evidence from anterograde tracing methods has shown that the projections of individual neurons are more diverse and divergent than previously indicated [62]. Second, in addition to common inputs, poly-synapotic inputs may explain these results. Because of the sampling rate of the recording and the slow kinetics of the Ca^2+^ sensor, we could not distinguish the correlations induced by mono-synaptic direct inputs from those induced by poly-synaptic direct inputs. Although the probability of mono-synaptic connections strongly depends on the distance between neurons, the probability of poly-synaptic connections may not. As correlations in Ca^2+^ imaging are combinations of correlations induced by both mono-synaptic and poly-synaptic inputs, it is possible to empirically observe spatially intermixed modules. Third, if recurrent synaptic connections are strong and variable across neurons, we would observe salt-and-pepper modules. In fact, recent studies utilizing both optogenetics and computational modeling have shown that salt-and-pepper responses to visual inputs in V1 can be explained by strong recurrent connections that are variable across neurons [63, 64].

To further clarify the underlying neural mechanisms of the module distribution, a direct causality analysis, instead of the correlation analysis employed in this study, is needed. For example, perturbative approaches, such as optogenetic stimulation during imaging, can help assess the existence of causal interactions through direct synaptic connections [65, 66]. This step towards elucidating causal relationships may offer deeper insights into the intricate mechanisms that shape spatially intermixed modules.

### Bridging the gap between macro and micro

The significance of our multi-scale analysis lies in revealing the spatial scales, from the single-cell scale to the mesoscale, at which changes in functional network structures commonly associated with the reduction of consciousness are observed. We found that the modularity measure could distinguish an awake state from sleep and anesthesia at the single-cell scale but not at the mesoscale (when the number of neighboring neurons *n*_*nei*_ *>* 10; Fig. 7c, e). This finding suggests that investigating functional network structures in relation to the reduction of consciousness requires examining not only the traditionally studied macro/mesoscale but also the single-cell scale. Our study serves as a starting point to inspire future research aimed at identifying the spatial scales that are fundamentally important for consciousness or, more generally, for brain information processing. Such investigations could be pursued experimentally [67–72] or through theoretical modeling and simulations [32, 73–75]. Even when studying mesoscale functional networks, it is important to note that the mesoscale data obtained through direct coarse-graining of single-neuron activity, as in our study, are qualitatively different from those obtained through non-invasive and indirect recording of neural activity, such as fMRI and EEG. In fact, previous studies using macro/mesoscale network analyses based on fMRI and EEG data have reported greater network segregation during sleep and anesthesia compared to an awake state [8–18], while this result was not observed in our mesoscale network analysis. These discrepancies are likely caused not only by the differences in direct versus indirect mesoscale recordings but also by other factors, including the spatial coverage of the recordings, temporal resolution. To further bridge the gap between micro- and mesoscale networks, the analysis of single-cell level resolution data, as conducted in this study, will be necessary.

Regarding the spatial distribution of modules at different spatial scales, our results support two seemingly opposing perspectives of brain network organization, namely the localistic and holistic views, depending on the activity resolution, i.e., macro/meso vs. micro. The former viewpoint posits that the brain is parceled into localized regions and that each region is responsible for specific information processing. For example, many functional network studies have reported that brain networks are segregated into spatially localized modules [17, 23, 30, 61, 68] that are thought to represent groups of nodes that perform similar brain functions and are responsible for specialized information processing [48]. This view is consistent with the theory of brain function localization [76, 77]. On the other hand, the latter viewpoint posits that cortical information processing is highly distributed; that is, the brain is not as neatly parceled into localized regions, as shown in the brain atlas. For example, we have previously reported that accurate somatosensory perception [78], perceptual memory consolidation [79], and memory enhancement [80] require coordinated activity in the primary somatosensory and secondary motor cortices. It has also been reported that behavior-related signals are represented brain-wide across species [81], including mice [82–84], flies [85], and worms [86]. In addition, some studies utilizing single-cell resolution recording techniques have reported that neurons with different stimulus selectivity are distributed in a spatially intermixed manner, such as in the visual cortex of rats [59] and in the sensory cortex of mice [60]. These results suggest that the brain functions in a more distributed manner by coordinating different regions as a whole, which is consistent with the holism of brain function viewpoint [81, 87]. Our single-cell level network analysis (Fig. 5) supports the holistic perspective while our coarse-grained network analysis (Fig. 7) supports the localistic one. Therefore, our results may provide new insights that can bridge the gap between the two perspectives: brain functions are macro/mesoscopically localized and microscopically intermixed in the cortex.

## Materials and Methods

### Resource availability

#### Lead Contact

Further information and requests for resources and reagents should be directed to and will be fulfilled by the Lead Contact, Masafumi Oizumi or Masanori Murayama.

#### Materials availability

All plasmids and AAVs generated in this study are available from the Lead Contact with a completed Materials Transfer Agreement.

### Data and code availability

Data and custom-written software used for data acquisition analysis can be requested from the lead contact.

### Experimental model and subject details

#### Animals

All animal experiments were performed in accordance with institutional guidelines and approved by the Animal Experiment Committee at RIKEN. Wild-type mice (C57BL/6JJmsSlc; Japan SLC, Shizuoka, Japan) were used. Male and female mice (12–24 weeks old) were used in this study. In all the experiments, the mice were housed in a 12 h-light/12 h-dark light cycle environment with *ad libitum* access to food and water.

### Method details

#### Adeno-associated virus (AAV) vector preparation

G-CaMP7.09 [88] was subcloned into the synapsin I (SynI)-expressing vector from the pN1-G-CaMP7.09 vector construct. The AAV AAV-DJ-Syn-G-CaMP7.09-WPRE was produced as previously described [89].

#### Neonatal AAV injection and surgery

Neonatal AAV injections and surgery for two-photon imaging were performed using previously described protocols [44, 45] with slight modifications to the surgical procedure.

Neonatal mice (postnatal day 0–2) were collected from their cages and anesthetized on ice (cryoanesthesia) for 2–3 min before injection, and then mounted in a neonatal mouse head holder (custom-made, NARISHIGE). The AAV (diluted to 4.0 *×*10^12^ GC/mL with *×*1 phosphate buffer saline (PBS), 4 µL per pup) was injected into the neonatal cortex using a glass pipette (Q100-30-15, Sutter Instrument) with a diameter of 45–50 µm. The injection site was located at a depth of 250–300 µm in the frontal area. The injection procedure was completed within 10 min after cryoanesthesia initiation, and the pups were warmed until their body temperature returned to normal. After the pups began to move, they were returned to their mothers.

For surgery, mice (8–20 weeks) were anesthetized with isoflurane (2%). Once the mice failed to react to stimuli, we administered hypodermic injections of the combination agent [90]—Medetomidine/Midazolam/Butorphanol (MMB)—at a dose of 5 mL/kg body weight for the procedure. The MMB solution consisted of medetomidine (0.12 mg/kg body weight), midazolam (0.32 mg/kg body weight), butorphanol (0.4 mg/kg body weight), and saline. After anesthesia with MMB, the head was shaved and placed in head holders (SG-4N, NARISHIGE). During surgery, their body temperatures were maintained at 36–37^*°*^C using a feedback-controlled heat pad (BWT-100, Bio Research Center), and their eyes were coated with an ointment (Neo-Medrol EE Ointment, Pfizer Inc.). After removing the scalp, a 5-mm diameter cranitomy was performed over an area that included the primary somatosensory area of the right hemisphere. The craniotomy was then covered with a 5-mm diameter No.1 cover glass (Matsunami Glass Ind.) and sealed with dental cement (Super Bond, Sun Medical). After cranial window implantation, EEG and EMG were performed as described in previous reports [79, 91, 92]. Briefly, an EEG screw was implanted over the V2L region in the right hemisphere and over the parietal cortex in the left hemisphere and locked with dental cement. Separate EEG signals were referenced to the screws located in the cerebellar cortex. For the EMG recording, a flexible wire cable was implanted into the neck muscle. A stainless-steel headplate (custom-made, ExPP Co., Ltd.) was then cemented on the skull over the cerebellum as parallel to the window glass as possible. The exposed skull, EEG screws, and EMG wires were covered with dental cement. After surgery, the mice were treated with a pesticidine-reversing agent, atipamezole hydrochloride (ANTISEDAN, Zoetis Inc.) solution at a dose of 0.12 mg/kg body weight. The mice were then placed on a heating pad to recover.

Skull images, including the cranial window of the head-fixed mouse, were acquired from one-photon macroscopic imaging after surgery to estimate the brain regions in the subsequent ROI analysis. The FOV was 13.3 mm*×* 13.3 mm and the image resolution was 1, 024 *×*1, 024 pixels after 2 *×*spatial binning. The excitation light intensity from a blue LED source (LEX2-B-S, Brain Vision Inc.) and the gain and exposure time of the sCMOS camera (Zyla 5.5, Andor) were tuned so that none of the pixels were saturated.

### In vivo two-photon calcium imaging of sleeping and anesthetized mice

In vivo two-photon imaging was performed using a custom-designed wide-field two-photon laser scanning microscope (FASHIO-2PM) [45]. To observe the spontaneous sleep-wake status in mice under the microscope, the mice were fixed in a custom-made stage box (ExPP Co., Ltd.) by firmly screwing the headplate to the stage, and then covered with an enclosure to maintain body temperature. The floor of the stage box was covered with the same bedding as that of the home cage. After recovery from surgery, the mice were head-fixed in a stage box daily during the light phase. The duration of head fixes increased daily from 1 min to 2 h in a steady manner to ensure that the head-fixed condition did not stress the mice. More than 2 weeks of habituation allowed the mice to show spontaneous wake/NREM/REM sleep states in the stage box. To observe neural activities during anesthesia, the mouse was anesthetized with 0.6% isoflurane inhalation after 20–60 min-wake recordings. Before each imaging session, the imaging window was cleaned with a cotton swab soaked in acetone. The implanted EEG/EMG electrodes were connected to the connector. Fluorescence was observed in the somas of layer 2/3 neurons (120–150 µm below the surface). G-CaMP7.09 was excited at 920 nm using a tunable Ti:Sa laser (Maitai DeepSee, Spectra Physics) and the laser power was set to 60–80 mW at the front of the objective lens. The fluorescence of G-CaMP7.09 was detected with a GaAsP PMT in the range of 515–565 nm. All imaging sessions were performed for approximately 3–4 h/day during the light phase (Zeitgeber time 3–7) for sleep imaging, or during the dark phase (Zeitgeber time 15–19) for anesthesia recordings at 7.65 frames/s with 2,048 × 2,048 pixels using custom-built software (Falcon, Nikon) and were saved as 16-bit monochrome tiff files.

### EEG and EMG recording and analysis

EEG and EMG signals were recorded during the imaging experiments. EEG and EMG signals were amplified and filtered (EEG, 0.1–100 Hz; EMG, 5–300 Hz) using an analog amplifier (MEG-6116, NIHON KOHDEN). The signals were digitized using a 16-bit analog-to-digital converter (Digidata 1440A, Molecular Devices) at a sampling rate of 1 kHz and acquired using Clampex software (v. 10.7, Molecular Devices). To temporally match the EEG/EMG data with two-photon images, the scan-timing signals for two-photon imaging were digitized using the same system. Post-hoc sleep state determination was performed as previously described [79, 93]. EEG signals from the same hemisphere as the imaging were used for post-hoc analysis. Briefly, the root-mean-square (RMS) of EMG signals, EEG delta (1–4 Hz) power (normalized to power in 1–50 Hz), and theta (6–9 Hz) power (normalized to power in 1–4 Hz) were calculated in each 4-s sliding window, which is nearly equal to 30 frames of two-photon imaging. We defined brain states as follows: an awake state, RMS value of EMG was above the threshold, which was defined manually in each animal; NREM, RMS value of EMG was below the threshold, and delta power/theta power was above 0.3; REM, the RMS value of the EMG was below the threshold, and the delta power/theta power was below 0.3. Sleep episodes of 12 s or less were dismissed and integrated into the sleep states before and after the episode. The transition from an awake state to REM was considered a quiet awake state.

### Imaging data analysis

All the obtained images were first subjected to motion correction using NoRMCorre [94]. We then used a low-computational-cost cell-detection (LCCD) algorithm [95] to identify the ROIs corresponding to single neurons. The fluorescence time course *F*_*ROI*_ (*t*) of each ROI was calculated by averaging all pixels within the ROI, and the fluorescence signal of the cell body was estimated as *F* (*t*) = *F*_*ROI*_ *− r× F*_*neuropil*_(*t*), where *r* = 0.7. The neuropil signal was defined as the average fluorescence intensity between 4 and 9 pixels from the boundary of the ROI (excluding all ROIs). The ΔF/F was calculated as (*F* (*t*) *−F*_0_(*t*))*/F*_0_(*t*), where *t* is time and *F*_0_(*t*) is the baseline fluorescence estimated as the 8% percentile value of the fluorescence distribution collected in a*±* 30 s window around each sample time point [96]. We further corrected the baseline with the part of “estimate baseline” function of Suite2P [97]. The SNR of each ROI’s ΔF/F was calculated using the “snr” function in MATLAB (MathWorks), where the signal was the ΔF/F below 0.5 Hz and the noise was the ΔF/F above 0.5 Hz. To exclude ROIs contaminated by noise, we selected ROIs that satisfied the following conditions: (1) The signal-to-noise ratio of ΔF/F was in the top 90%, (2) ROIs not obscured by nearby blood vessels, (3) the range of ΔF/F values fell between *−* 1 and 10, (4) ROIs with at least one Ca^2+^ transient during recording, (5) the neuropil fluorescence was lower than that of the soma, (6) less than 45% of the frames had a ΔF/F lower than average, and (7) the correlation between ΔF/F and the square root of the EMG values was less than 0.2. See Tables S1 and S2 for the number of ROIs after excluding contaminated ones. We then deconvolved ΔF/F signals using OASIS [98]. Subsequently, we smoothed the deconvolved signals by applying a Gaussian filter with a length of 1.96 seconds (15 frames).

All analyses to detect blood vessels were performed semi-automatically using ImageJ software [99] as follows. The median image of the two-photon images for 500 frames was converted into the frequency domain via a 2D Fourier transform. After extracting the low-frequency components of the image, an inverse Fourier transform was performed. Then, the blood vessel was detected by thresholding methods plugged in with ImageJ as function “AutoThreshold.” The algorithm “MinError” was used for this.

### Estimation of Functional Networks

To estimate the functional networks, we calculated the Pearson correlation coefficients of the deconvolved Ca^2+^ signal between the neuron pairs. Only neurons that were not silent in any of the windows were included in the analysis (see Tables. S1 and S2 for the number of neurons used for network analysis). The window length was 196 s (1500 frames) for the wakefulness-sleep data and 379 s (2900 frames) for the wakefulness-anesthesia data. These window lengths are the longest duration feasible for analysis across all individuals. For the wakefulness-sleep data, since the sleep state of a mouse typically does not persist for more than 196 s, we segmented and combined the contiguous sections of equivalent states to achieve a cumulative duration of 196 s.

We excluded time frames that could introduce artifacts or ambiguity. We excluded frames with EMG potentials above a threshold in the previous and following 6.5 s (50 frames) to avoid motion-induced artifacts. We also excluded frames in which the same state (either an awake state or NREM sleep) did not persist for more than 39 s (300 frames), to avoid ambiguity in determining whether the mouse was awake or asleep. For the anesthesia data, we excluded the first 1000 frames after the start of anesthesia to avoid the possibility that the state is not yet stable.

We binarized the correlation matrices at a threshold, setting it such that the ratio of the number of edges to all possible node pairs was equal to *K* [49]. For example, if *K* = 0.01, neuron pairs whose absolute correlation values rank in the top 1 percent are considered to have an edge.

### Computation of confidence interval for the difference between an awake state and NREM sleep or between an awake state and anesthesia

To assess the statistically likely ranges of the differences in network measures between an awake state and NREM sleep for each connection density *K*, confidence intervals were calculated for the difference. As the network measures were estimated multiple times using different time windows for each state within each mouse, the network measures were first averaged for each mouse and state. The average network measures of NREM sleep or anesthesia were subtracted from those of an awake state for each mouse. Subsequently, the 95% confidence interval of the difference in network measures between an awake state and NREM sleep or between an awake state and anesthesia was computed based on a one-sample *t*-test. For the wakefulness-sleep data, sample sizes were 5 with 4 degrees of freedom. For the wakefulness-anesthesia data, sample sizes were 4 with 3 degrees of freedom.

### Graph Metrics

We calculated graph-theoretical measures using the Brain Connectivity Toolbox [6] and custom-made MATLAB codes.

### Modularity

Modularity is defined as the number of edges within modules minus the expected number of such edges [100],

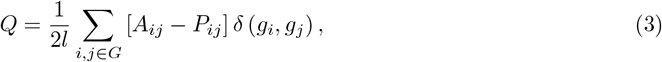

where *G* is the set of all nodes in the network, *l* is the number of edges in the network, *A*_*ij*_ is an element of the adjacency matrix,

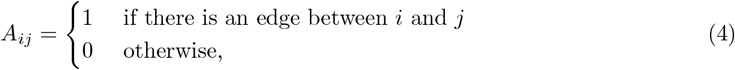

*P*_*ij*_ is the expected number of edges between *i* and *j*,

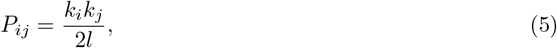

*k*_*i*_ is the degree of a node *i, g*_*i*_ is the module to which node *i* belongs, and *δ*(*r, s*) is defined as

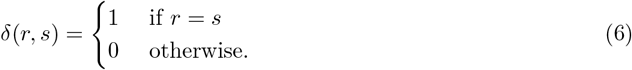

We identified a module assignment using the Louvain algorithm [101], which divides a network into modules to maximize *Q*, and then calculated the modularity. Because the Louvain algorithm is stochastic, we executed it 10 times for each network and chose a module assignment where the value of modularity was the largest. In the Louvain algorithm, the number of modules is automatically determined to maximize the modularity value. When we compared the modularity values by setting the number of modules, as shown in Figs. S9 and S10, we set it to the smallest number of modules in the network that we wanted to compare. Iteration used the following process: selecting the two modules that resulted in the least decrease in modularity and then merging them.

We introduce a node-specific modularity measure, *Q*_*i*_, which quantifies the extent to which a node

*i* contributes to the modularity *Q*,

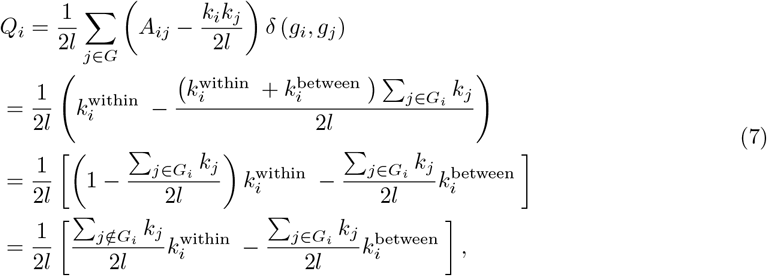

where *G*_*i*_ is the set of nodes that belong to the same module as node *i*,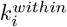is the within-module degree of a node *i*, and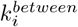 is the across-module degree of a node *i*. The last line in Eq. (7) shows that *Q*_*i*_ is large when node *i* has many within-module connections, i.e.,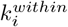 is large, and few across-module connections, i.e.,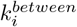 is small. Comparing eq. (3) and the first line in eq. (7), we can see that *Q*_*i*_ is the exact decomposition of the modularity index *Q*,

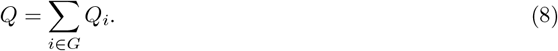

Although participation coefficient and within-module degree z-score are the most commonly used measures for identifying connector nodes and provincial nodes [102, 103], these measures are not directly related to the modularity of the network. On the other hand, *Q*_*i*_ is the exact decomposition of the modularity index *Q*, thus enabling us to precisely characterize the specific contributions of individual neurons to the overall modular structure of the network.

### Global efficiency and transitivity

We also calculated the global efficiency and the transitivity as measures of network integration and segregation. The global efficiency is defined as the average inverse shortest path length,

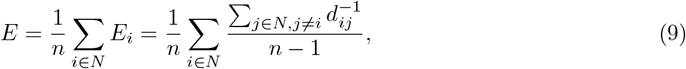

where *N* is the set of all nodes in the network, *n* is the number of nodes, and *d*_*ij*_ is the shortest path length between node *i* and *j* [104]. This measure quantifies how efficiently one node can communicate with another. The transitivity is defined as the fraction of triangles in a network,

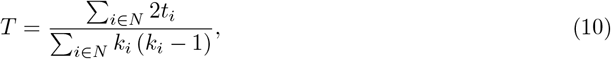

where *t*_*i*_ is the number of triangles around node *i* and *k*_*i*_ is the degree of node *i* [105]. A large number of triangles implies segregation.

### Declaration of generative AI and AI-assisted technologies in the writing process

During the preparation of this work the authors used ChatGPT in order to improve language and readability. After using this tool, the authors reviewed and edited the content as needed and take full responsibility for the content of the publication.

## Acknowledgements

We thank Muneki Ikeda for valuable comments and discussions. This research was supported by the Japan Society for the Promotion of Science (JSPS) KAKENHI Grant Numbers JP19J01973 to I.O., JP20H05775 to M.M., 20H05712 and 23H04834 to M.O., the AMED-Brain/Minds Project Grant Number JP15dm0207001 to M.M., KAO Corp. to M.M., the Toray Science Foundation to M.M., Japan Science and Technology Agency (JST) CREST Grant Number JPMJCR1864 to D.K., J.K., and M.O., JST SPRING Grant Number JPMJSP2108 to D.K., and JSPS KAKENHI Grant number 24KJ0762 to D.K..

## Supplementary Information

### Supplementary Tables

**Table S1:** Summary of basic information of wakefulness-sleep data

| Mouse ID | mouse 1 | mouse 2 | mouse 3 | mouse 4<br>(day 1) | mouse 4<br>(day 2) | mouse 5 |
| --- | --- | --- | --- | --- | --- | --- |
| # of originally detected ROIs | 10113 | 12639 | 11674 | 17601 | 19635 | 11796 |
| # of ROIs after excluding contaminated ones | 7843 | 6574 | 7112 | 9612 | 10197 | 7355 |
| # of ROIs used for the network analysis | 7693 | 6169 | 6629 | 8795 | 9487 | 6920 |
| # of sets of 196-second windows<br>used for network analysis (Awake) | 1 | 4 | 1 | 1 | 6 | 6 |
| # of sets of 196-second windows<br>used for network analysis (NREM) | 3 | 5 | 8 | 7 | 4 | 3 |
| Total duration of an awake state (sec) | 320 | 1133 | 471 | 578 | 2052 | 1411 |
| Total duration of NREM sleep (sec) | 732 | 1230 | 1743 | 1627 | 973 | 997 |
| Total duration of REM sleep (sec) | 72 | 0 | 123 | 235 | 42 | 51 |
| Total duration of a quiet awake state (sec) | 98 | 81 | 107 | 4 | 71 | 25 |

**Table S2:** Summary of basic information of wakefulness-anesthesia data

| Mouse ID | mouse 3 | mouse 5 | mouse 6 | mouse 7 |
| --- | --- | --- | --- | --- |
| # of originally detected ROIs | 9057 | 7571 | 9059 | 10400 |
| # of ROIs after excluding contaminated ones | 7350 | 6592 | 5295 | 4337 |
| # of ROIs used for the network analysis | 1900 | 3210 | 2824 | 3013 |
| # of sets of 379-second windows<br>used for network analysis (Awake) | 3 | 5 | 3 | 3 |
| # of sets of 379-second windows<br>used for network analysis (Anesthesia) | 2 | 1 | 4 | 2 |
| Total duration of an awake state (sec) | 1242 | 2043 | 1242 | 1242 |
| Total duration of anesthesia (sec) | 1242 | 514 | 1960 | 1242 |

### Supplementary Figures

**Figure S1:**
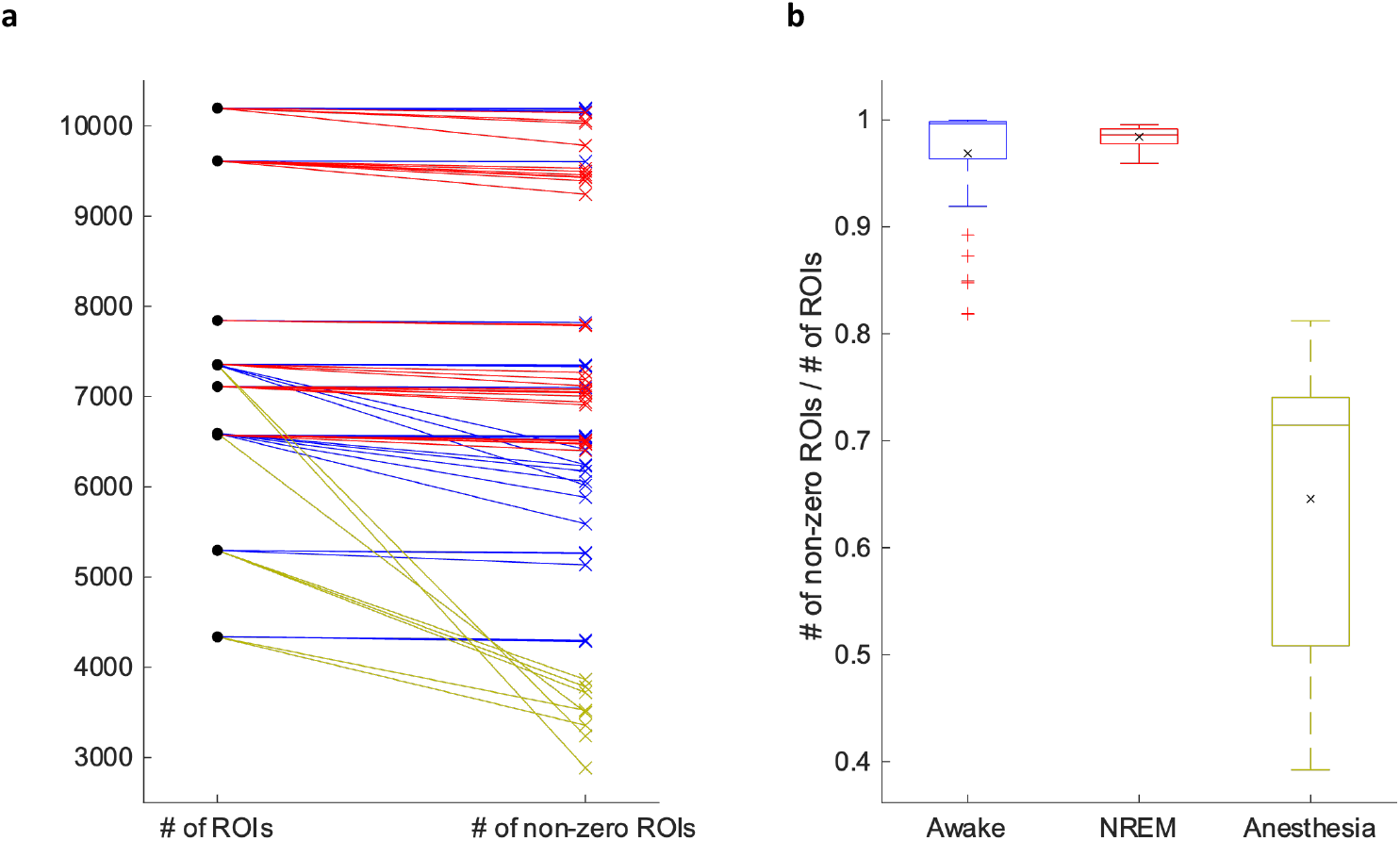
Number of neurons active within each time window. **a**. Number of neurons before (black dots) and after (colored crosses) excluding neurons with no Ca^2+^ event within each time window during an awake state (blue), NREM sleep (red), and anesthesia (yellow). Each black dot represents the number of neurons after excluding contaminated ones but before excluding ones with no Ca^2+^ event (see also Tables S1 and S2). Each colored cross corresponds to a single time window. The length of the time window is equal to that used in the subsequent network analysis (196s for the wakefulness-sleep data and 379s for the wakefulness-anesthesia data). Data of the same mice are connected by lines. **b**. Boxplot showing the ratio of neuron counts before and after excluding contaminated neurons. The black cross represents the average across time windows.

**Figure S2:**
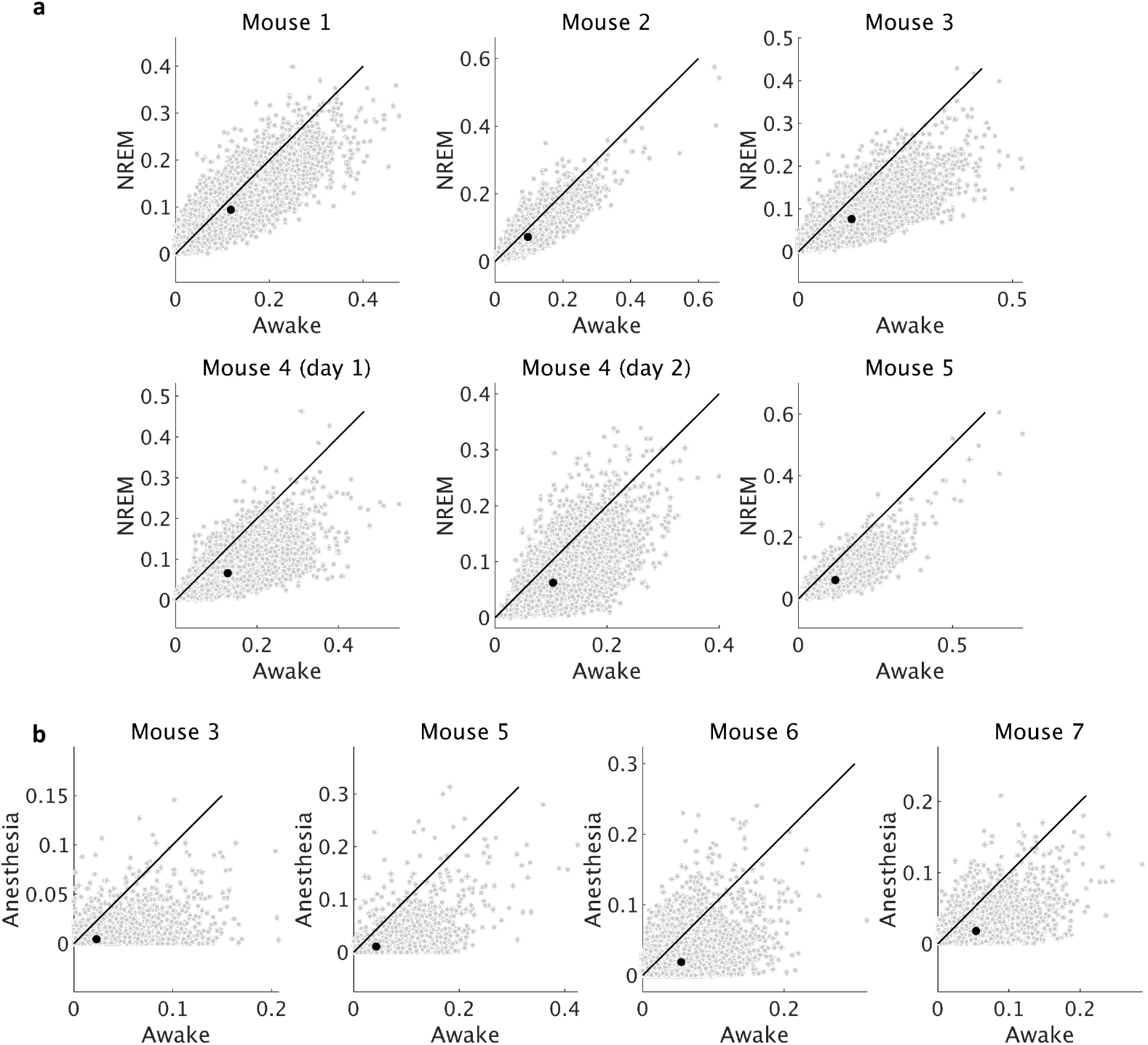
Distribution of event frequency of deconvolved signal of the wakefulness-sleep data (**a**) and the wakefulness-anesthesia data (**b**). The gray dots indicate individual neurons. The black dot indicates average values.

**Figure S3:**
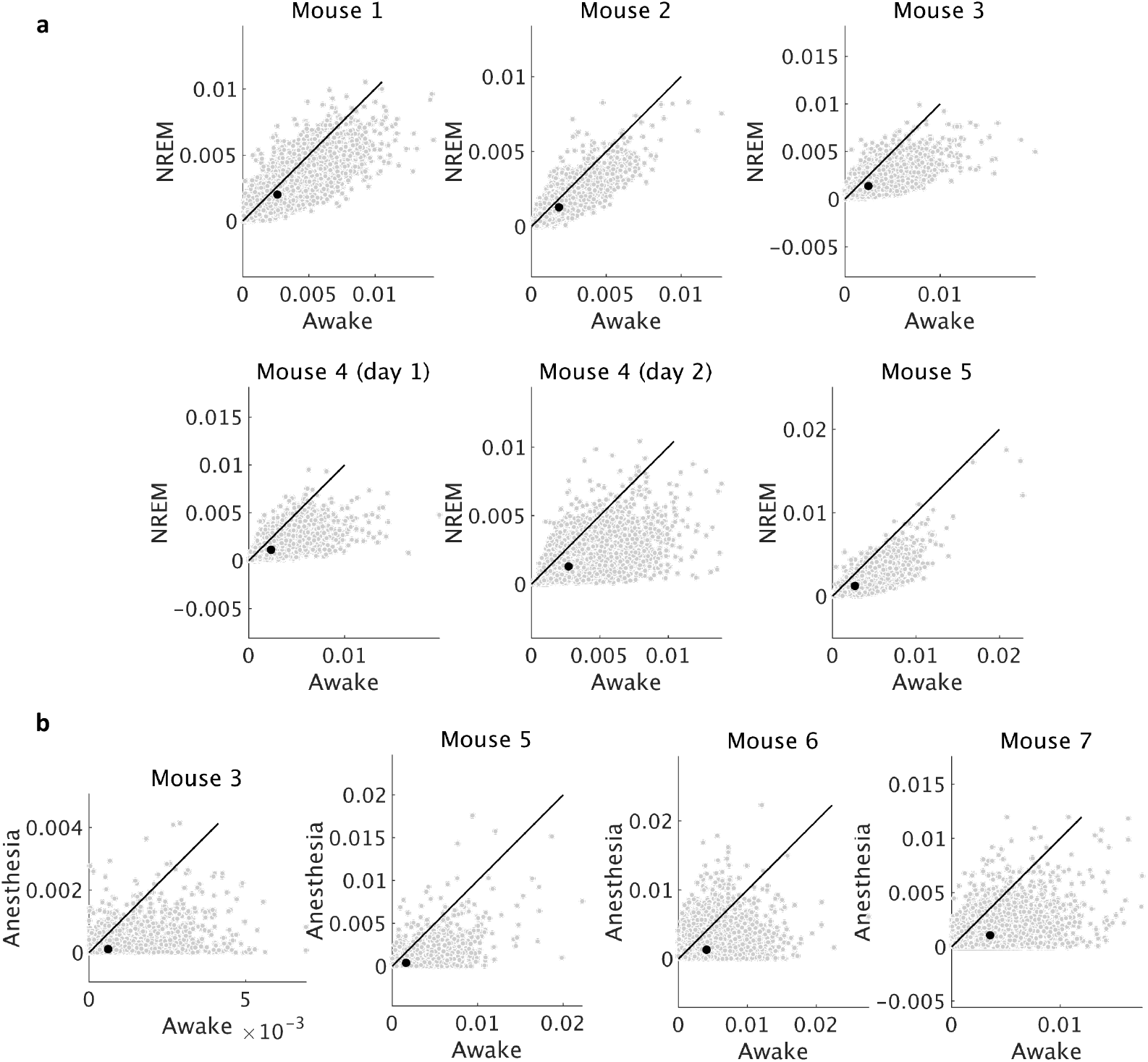
Distribution of mean amplitude of deconvolved signal of the wakefulness-sleep data (**a**) and the wakefulness-anesthesia data (**b**). The gray dots indicate individual neurons. The black dot indicates average values.

**Figure S4:**
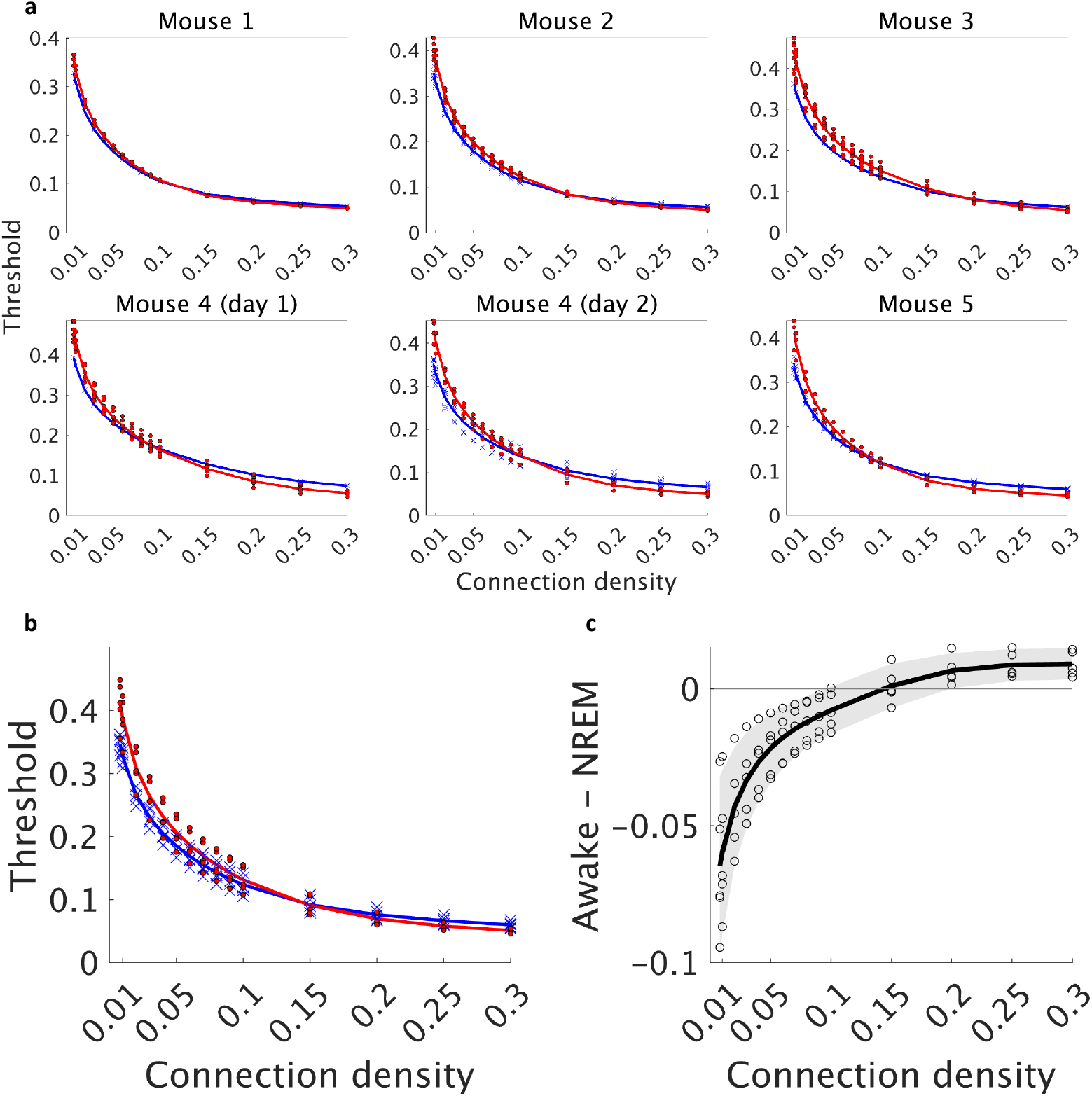
Dependence of the absolute correlation threshold on the connection density during an awake state and NREM sleep. **a**. Comparison of the threshold between an awake state and NREM sleep for each mouse. Each blue cross represents the threshold for a network during an awake state, while each red circle represents the threshold during NREM sleep. The blue and red solid lines represent the averages of the threshold for an awake state and NREM sleep, respectively. **b**. Comparison of the average threshold between an awake state (blue) and NREM sleep (red). Crosses (an awake state) and dots (NREM sleep) represent the threshold of individual mouse, averaged across networks estimated from different time windows. Lines represent the mean threshold across mice. **c**. Difference in the average threshold between an awake state and NREM sleep. Each circle represents an individual mouse. The black line represents the average across mice. The gray-shaded area indicates the 95% confidence interval (based on one-sample t-test, *n* = 5, *df* = 4).

**Figure S5:**
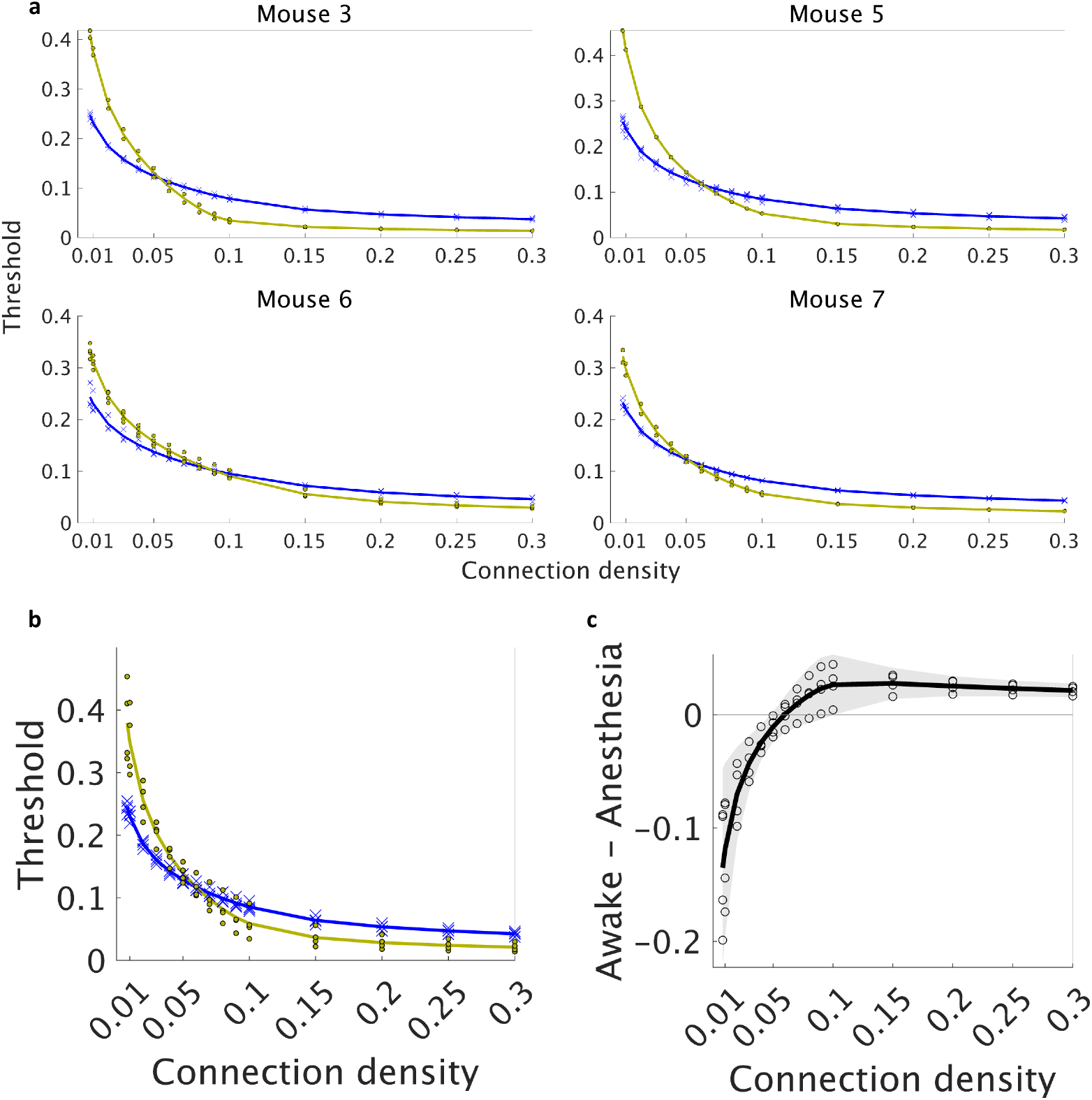
Dependence of the absolute correlation threshold on the connection density during an awake state and anesthesia. **a**. Comparison of the threshold between an awake state and anesthesia for each mouse. Each blue cross represents the threshold for a network during an awake state, while each yellow circle represents the threshold during anesthesia. The blue and yellow solid lines represent the averages of the threshold for an awake state and anesthesia, respectively. **b**. Comparison of the average threshold between an awake state (blue) and anesthesia (yellow). Crosses (an awake state) and dots (isoflurane-induced anesthesia) represent the threshold of individual mouse, averaged across networks estimated from different time windows. Lines represent the mean threshold across mice. **c**. Difference in the average threshold between an awake state and anesthesia. Each circle represents an individual mouse. The black line represents the average across mice. The gray-shaded area indicates the 95% confidence interval (based on one-sample t-test, *n* = 4, *df* = 3).

**Figure S6:**
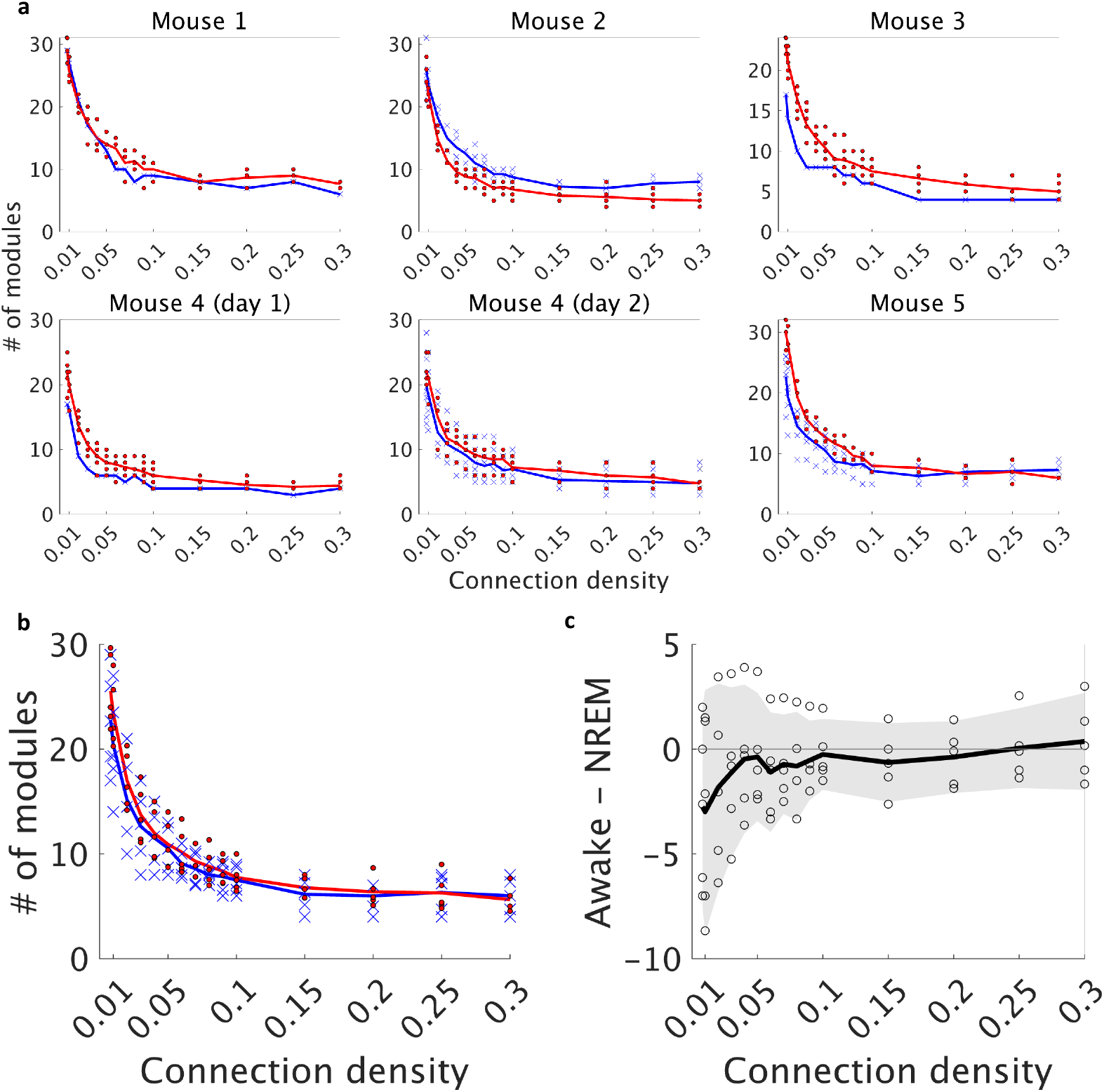
Comparison of the number of modules between an awake state and NREM sleep. **a**. Comparison for each mouse. Each blue cross represents the number of modules for a network during an awake state, while each red circle represents the number of modules during NREM sleep. The blue and red solid lines represent the averages of the number of modules for an awake state and NREM sleep, respectively. **b**. Comparison of the average number of modules between an awake state (blue) and NREM sleep (red). Crosses (an awake state) and dots (NREM sleep) represent the number of modules of individual mouse, averaged across networks estimated from different time windows. Lines represent the mean number of modules across mice. **c**. Difference in the average number of modules between an awake state and NREM sleep. Each circle represents an individual mouse. The black line represents the average across mice. The gray-shaded area indicates the 95% confidence interval (based on one-sample t-test, *n* = 5, *df* = 4).

**Figure S7:**
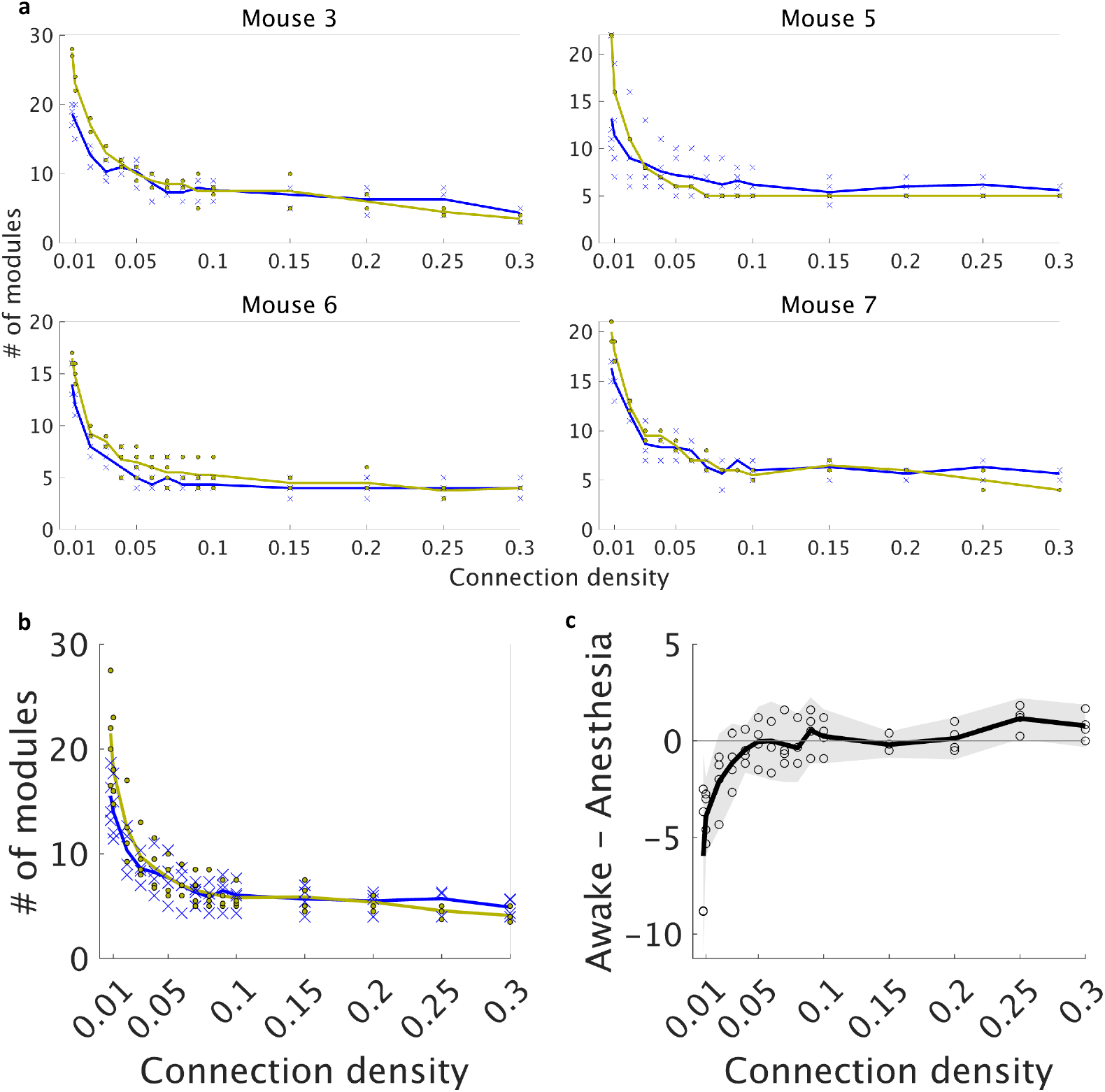
Comparison of the number of modules between an awake state and anesthesia. **a**. Comparison for each mouse. Each blue cross represents the number of modules for a network during an awake state, while each yellow circle represents the number of modules during anesthesia. The blue and yellow solid lines represent the averages of the number of modules for an awake state and anesthesia, respectively. **b**. Comparison of the average number of modules between an awake state (blue) and anesthesia (yellow). Crosses (an awake state) and dots (isoflurane-induced anesthesia) represent the number of modules of individual mouse, averaged across networks estimated from different time windows. Lines represent the mean number of modules across mice. **c**. Difference in the average number of modules between an awake state and anesthesia. Each circle represents an individual mouse. The black line represents the average across mice. The gray-shaded area indicates the 95% confidence interval (based on one-sample t-test, *n* = 4, *df* = 3).

**Figure S8:**
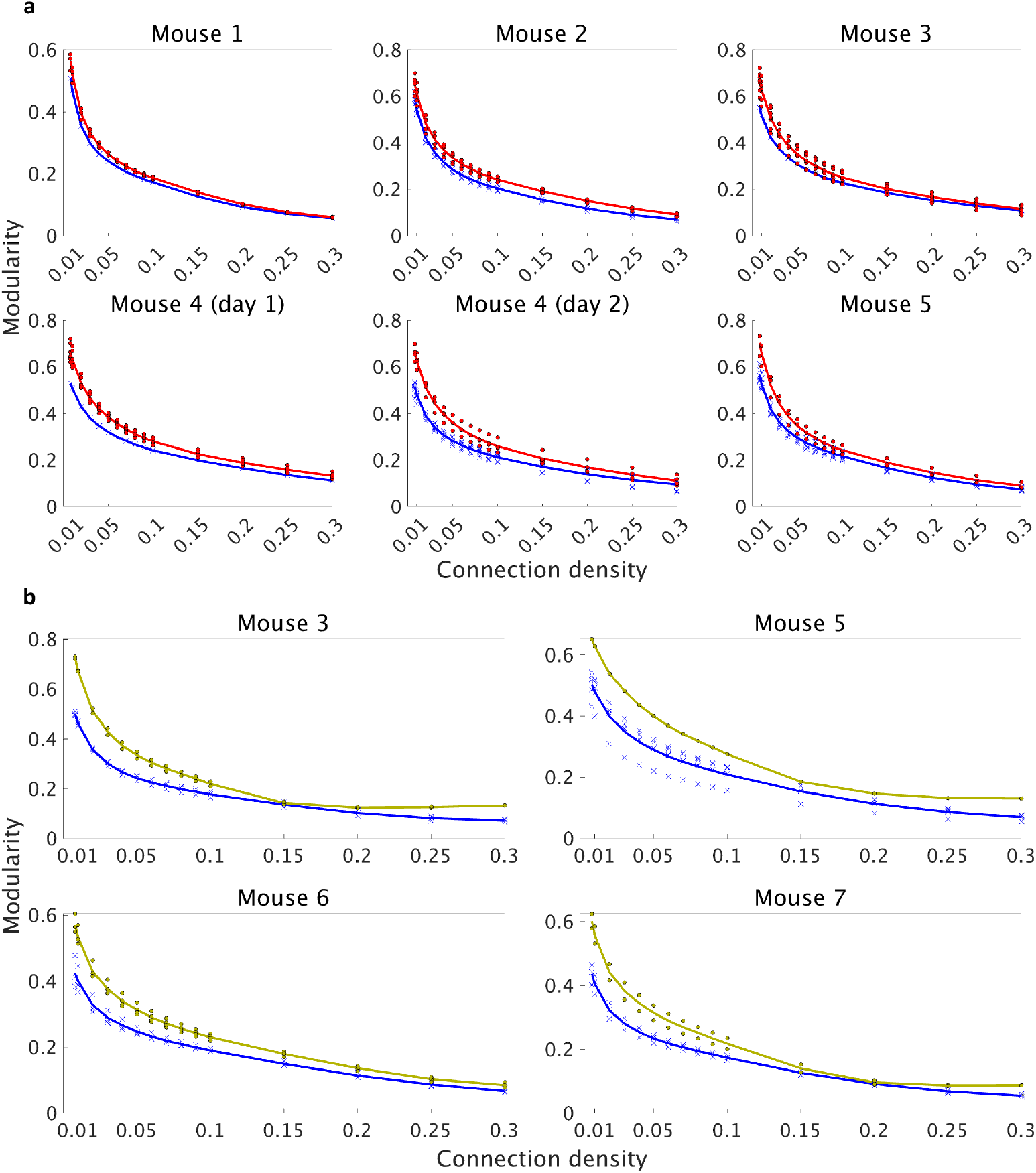
Comparison of modularity for each mouse. **a**. Each blue cross represents modularity for a network during an awake state, while each red circle represents modularity during NREM sleep. The blue and red solid lines represent the averages of modularity for an awake state and NREM sleep, respectively. **b**. Each blue cross represents modularity for a network during an awake state, while each yellow circle represents modularity during anesthesia. The blue and yellow solid lines represent the averages of modularity for an awake state and NREM sleep, respectively.

**Figure S9:**
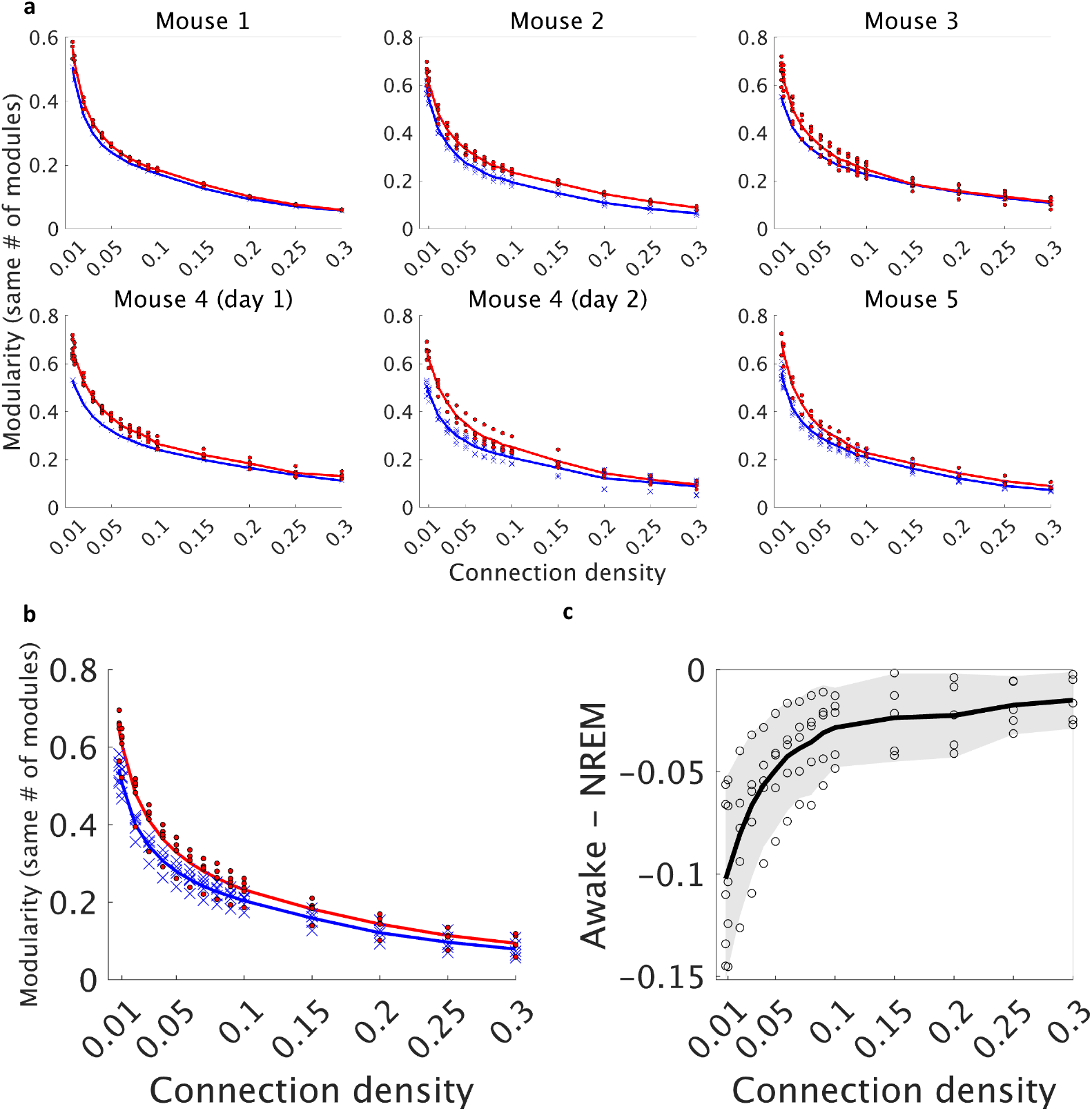
Comparison of modularity when the number of modules is matched between an awake state and NREM sleep. **a**. Comparison for each mouse. Each blue cross represents modularity for a network during an awake state, while each red circle represents modularity during NREM sleep. The blue and red solid lines represent the averages of modularity for an awake state and NREM sleep, respectively. **b**. Comparison of the average modularity between an awake state (blue) and NREM sleep (red). Crosses (an awake state) and dots (NREM sleep) represent modularity of individual mouse, averaged across networks estimated from different time windows. Lines represent the mean modularity across mice. **c**. Difference in the average modularity between an awake state and NREM sleep. Each circle represents an individual mouse. The black line represents the average across mice. The gray-shaded area indicates the 95% confidence interval (based on one-sample t-test, *n* = 5, *df* = 4).

**Figure S10:**
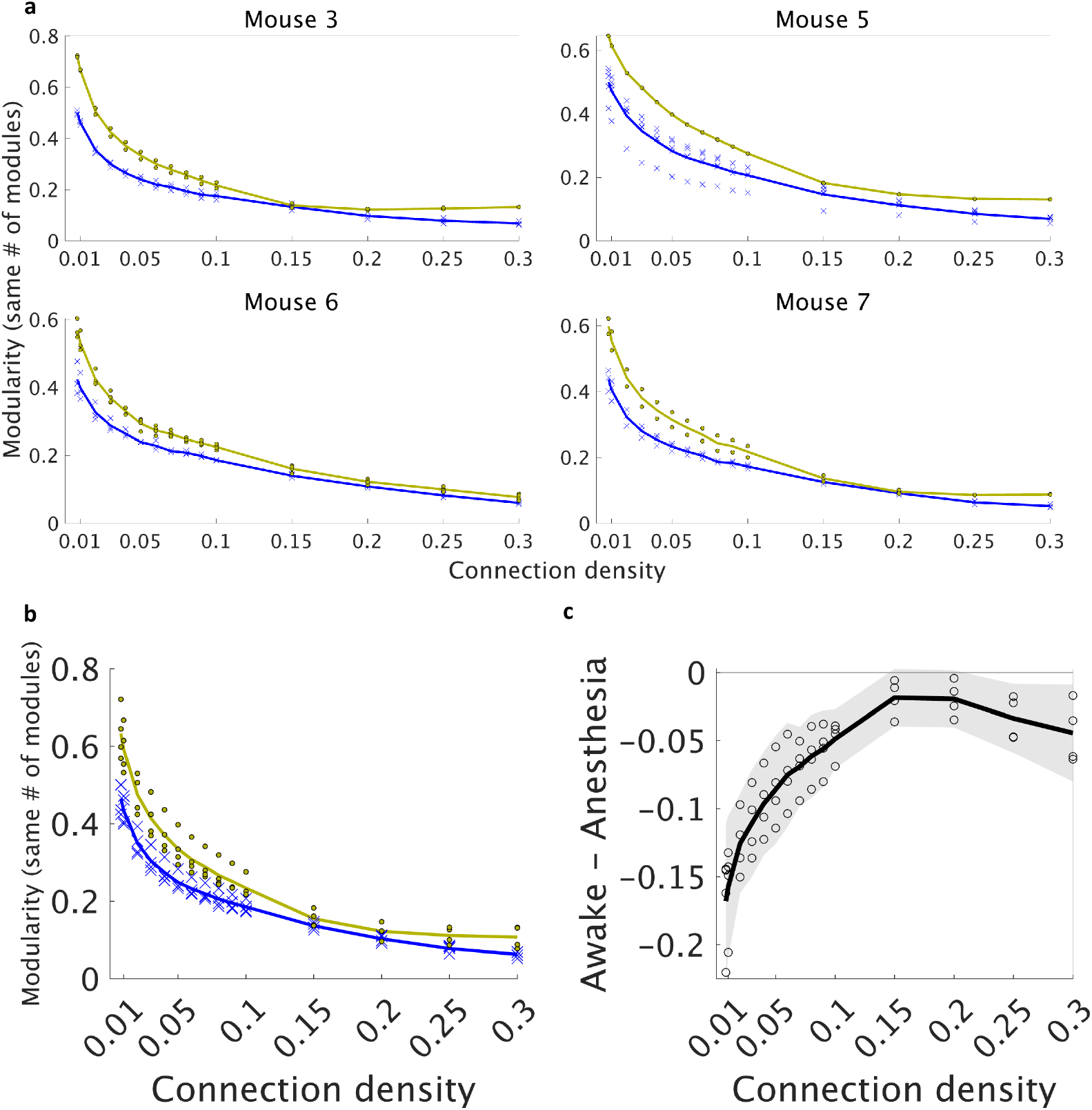
Comparison of modularity when the number of modules is matched between an awake state and anesthesia. **a**. Comparison for each mouse. Each blue cross represents modularity for a network during an awake state, while each yellow circle represents modularity during anesthesia. The blue and yellow solid lines represent the averages of modularity for an awake state and anesthesia, respectively. **b**. Comparison of the average modularity between an awake state (blue) and anesthesia (yellow). Crosses (an awake state) and dots (isoflurane-induced anesthesia) represent modularity of individual mouse, averaged across networks estimated from different time windows. Lines represent the mean modularity across mice. **c**. Difference in the average modularity between an awake state and anesthesia. Each circle represents an individual mouse. The black line represents the average across mice. The gray-shaded area indicates the 95% confidence interval (based on one-sample t-test, *n* = 4, *df* = 3).

**Figure S11:**
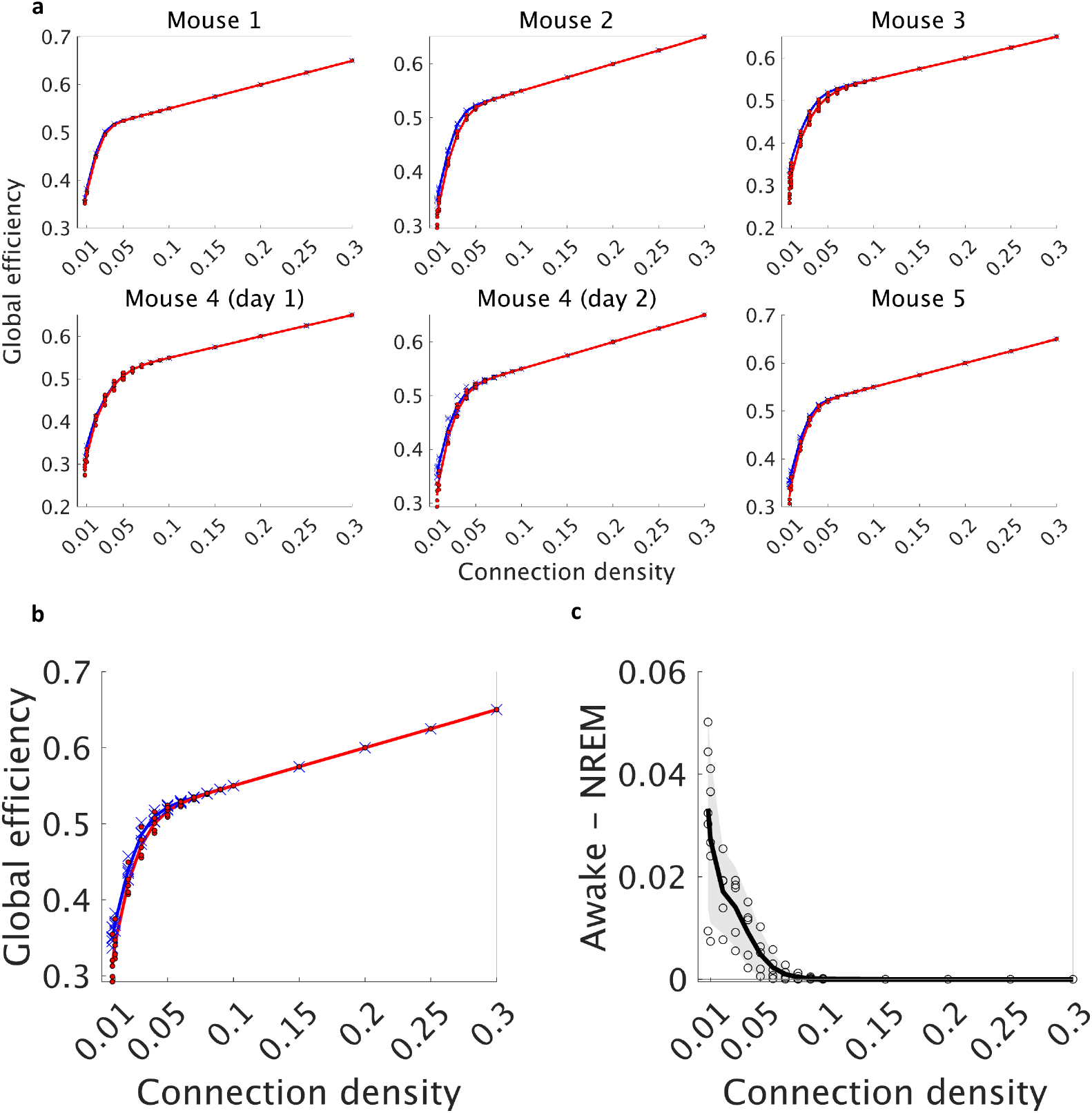
Comparison of global efficiency between an awake state and NREM sleep. **a**. Comparison for each mouse. Each blue cross represents global efficiency for a network during an awake state, while each red circle represents global efficiency during NREM sleep. The blue and red solid lines represent the averages of global efficiency for an awake state and NREM sleep, respectively. **b**. Comparison of the average global efficiency between an awake state (blue) and NREM sleep (red). Crosses (an awake state) and dots (NREM sleep) represent global efficiency of individual mouse, averaged across networks estimated from different time windows. Lines represent the mean global efficiency across mice. **c**. Difference in the average global efficiency between an awake state and NREM sleep. Each circle represents an individual mouse. The black line represents the average across mice. The grayshaded area indicates the 95% confidence interval (based on one-sample t-test, *n* = 5, *df* = 4).

**Figure S12:**
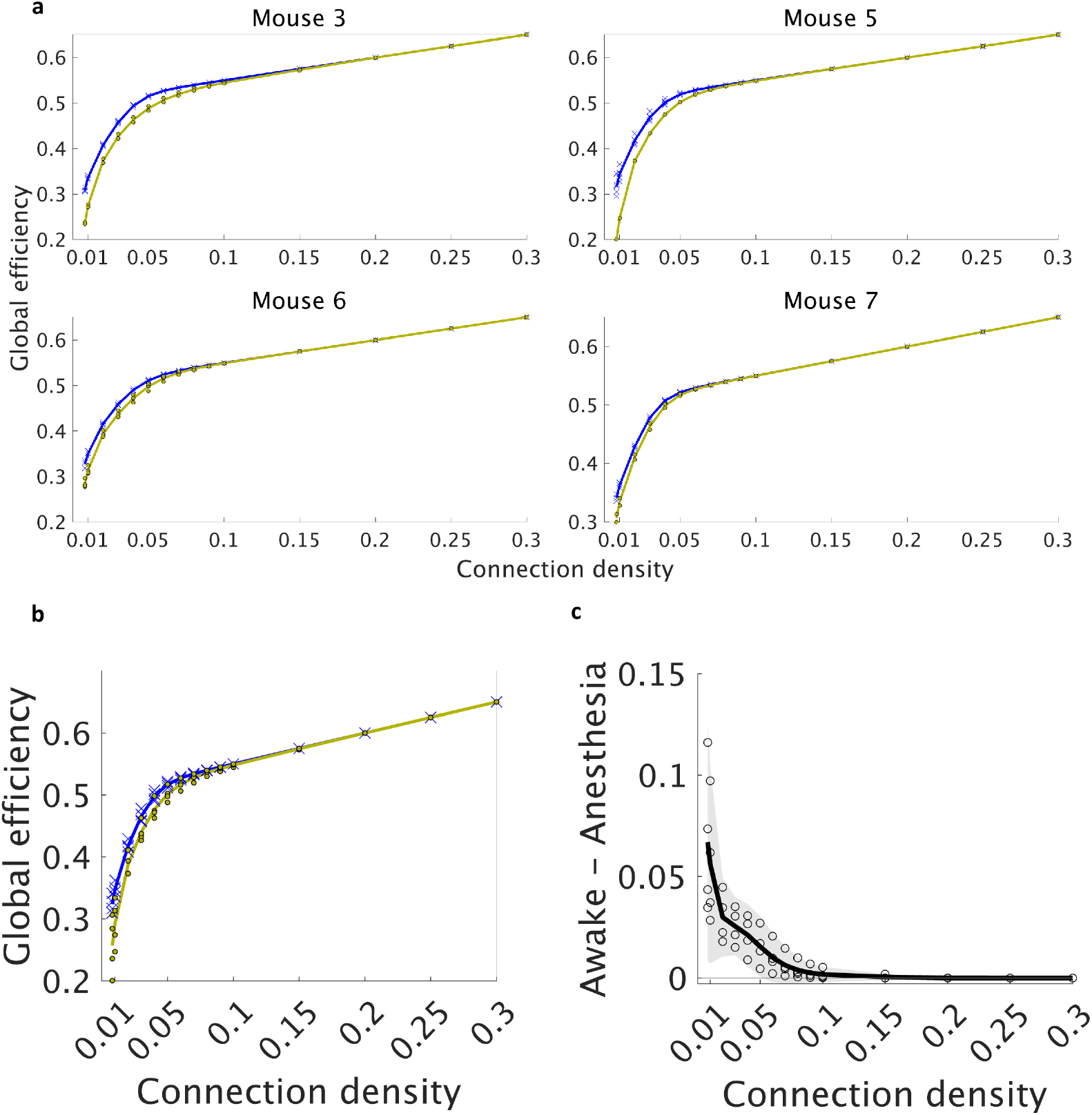
Comparison of global efficiency between an awake state and anesthesia. **a**. Comparison for each mouse. Each blue cross represents global efficiency for a network during an awake state, while each yellow circle represents global efficiency during anesthesia. The blue and yellow solid lines represent the averages of global efficiency for an awake state and anesthesia, respectively. **b**. Comparison of the average global efficiency between an awake state (blue) and anesthesia (yellow). Crosses (an awake state) and dots (isoflurane-induced anesthesia) represent global efficiency of individual mouse, averaged across networks estimated from different time windows. Lines represent the mean global efficiency across mice. **c**. Difference in the average global efficiency between an awake state and anesthesia. Each circle represents an individual mouse. The black line represents the average across mice. The gray-shaded area indicates the 95% confidence interval (based on one-sample t-test, *n* = 4, *df* = 3).

**Figure S13:**
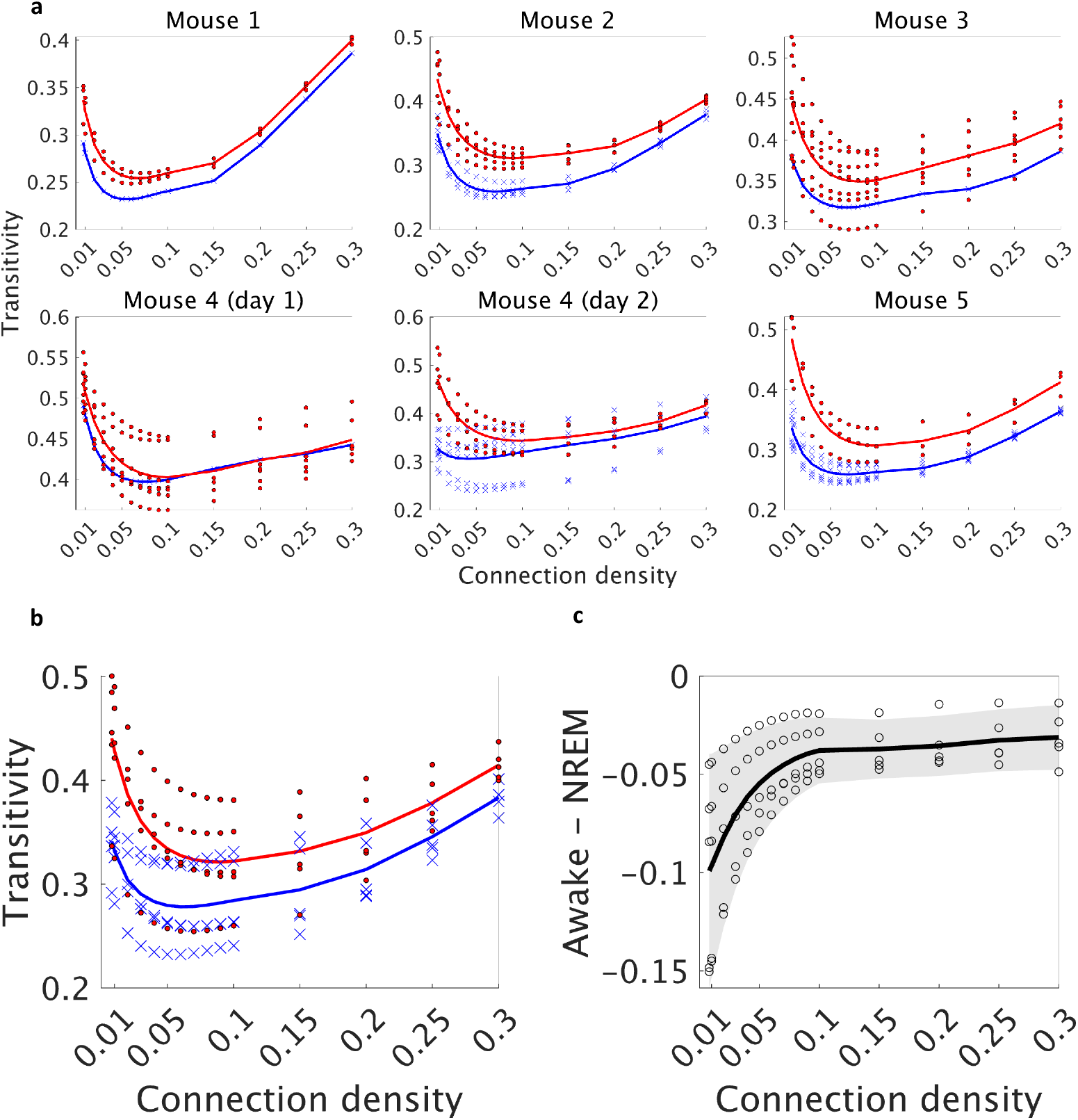
Comparison of transitivity between an awake state and NREM sleep. **a**. Comparison for each mouse. Each blue cross represents transitivity for a network during an awake state, while each red circle represents transitivity during NREM sleep. The blue and red solid lines represent the averages of transitivity for an awake state and NREM sleep, respectively. **b**. Comparison of the average transitivity between an awake state (blue) and NREM sleep (red). Crosses (an awake state) and dots (NREM sleep) represent transitivity of individual mouse, averaged across networks estimated from different time windows. Lines represent the mean transitivity across mice. **c**. Difference in the average transitivity between an awake state and NREM sleep. Each circle represents an individual mouse. The black line represents the average across mice. The gray-shaded area indicates the 95% confidence interval (based on one-sample t-test, *n* = 5, *df* = 4).

**Figure S14:**
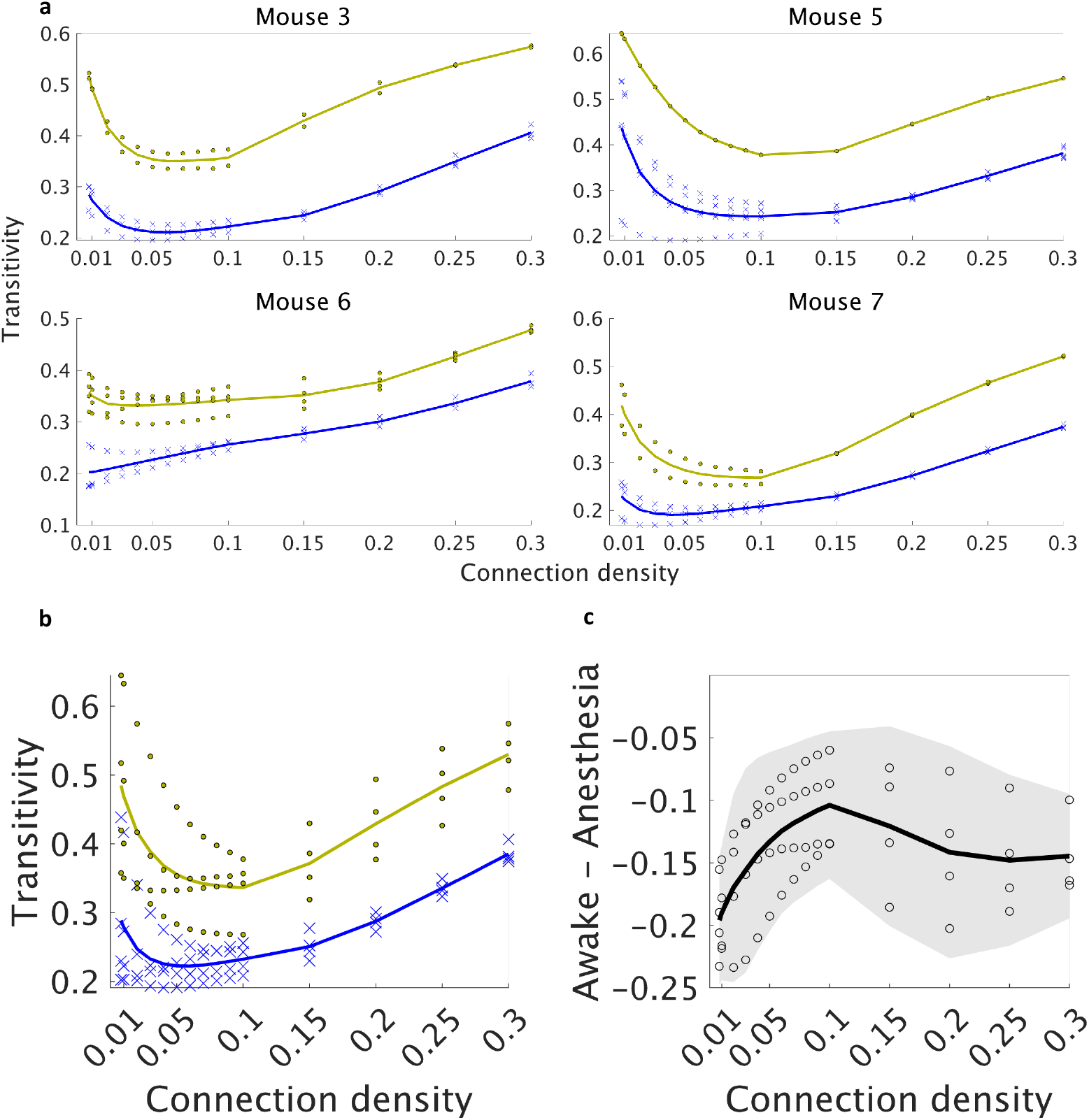
Comparison of transitivity between an awake state and anesthesia. **a**. Comparison for each mouse. Each blue cross represents transitivity for a network during an awake state, while each yellow circle represents transitivity during anesthesia. The blue and yellow solid lines represent the averages of transitivity for an awake state and anesthesia, respectively. **b**. Comparison of the average transitivity between an awake state (blue) and anesthesia (yellow). Crosses (an awake state) and dots (isoflurane-induced anesthesia) represent transitivity of individual mouse, averaged across networks estimated from different time windows. Lines represent the mean transitivity across mice. **c**. Difference in the average transitivity between an awake state and anesthesia. Each circle represents an individual mouse. The black line represents the average across mice. The gray-shaded area indicates the 95% confidence interval (based on one-sample t-test, *n* = 4, *df* = 3).

**Figure S15:**
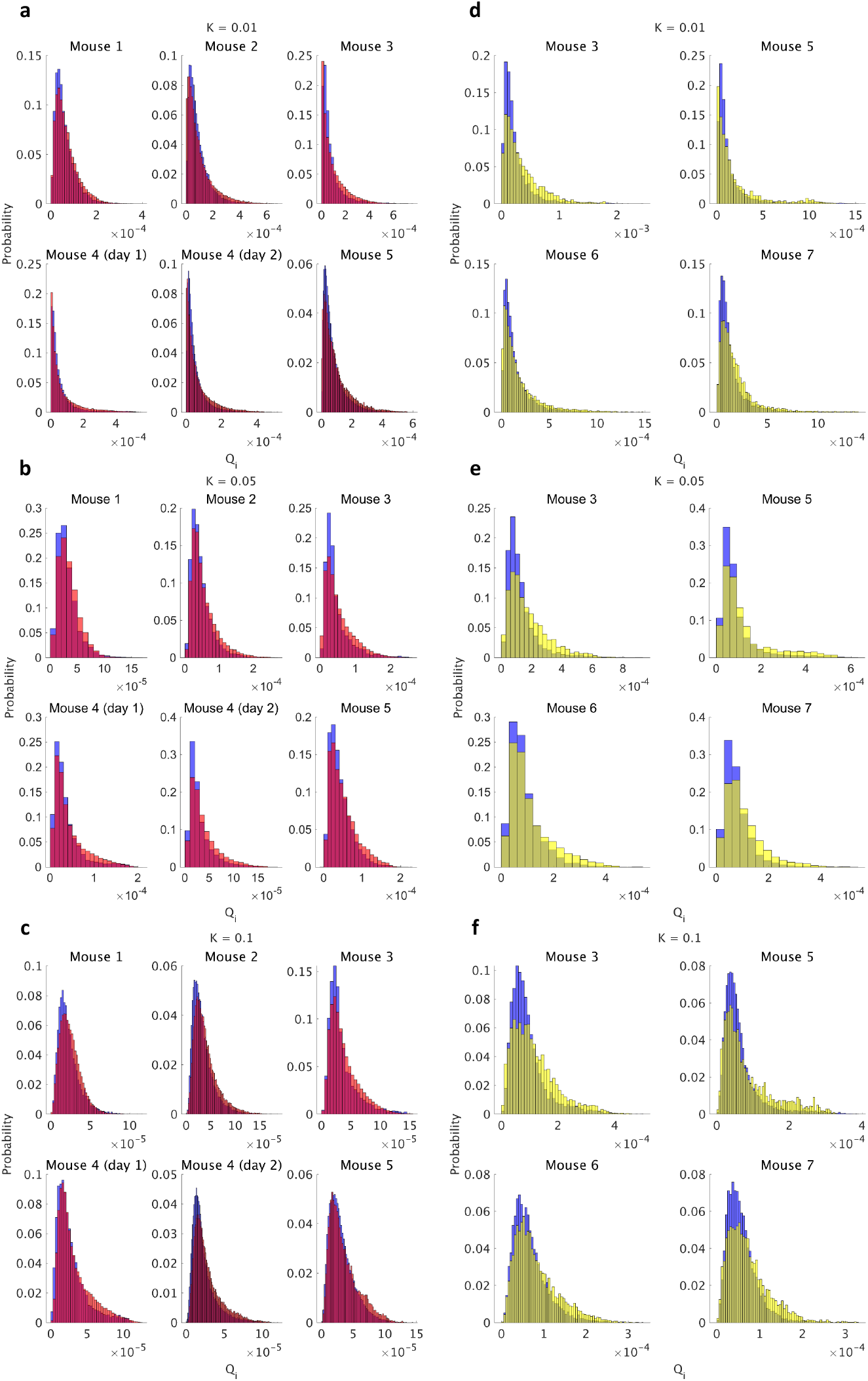
Histogram of *Q*_*i*_ during an awake state (blue) and NREM sleep (red) (**a–c**) as well as during an awake state (blue) and anesthesia (**d–f**) for different connection densities (**a, d** for *K* = 0.01; **b, e** for *K* = 0.05; **c, f** for *K* = 0.1).

**Figure S16:**
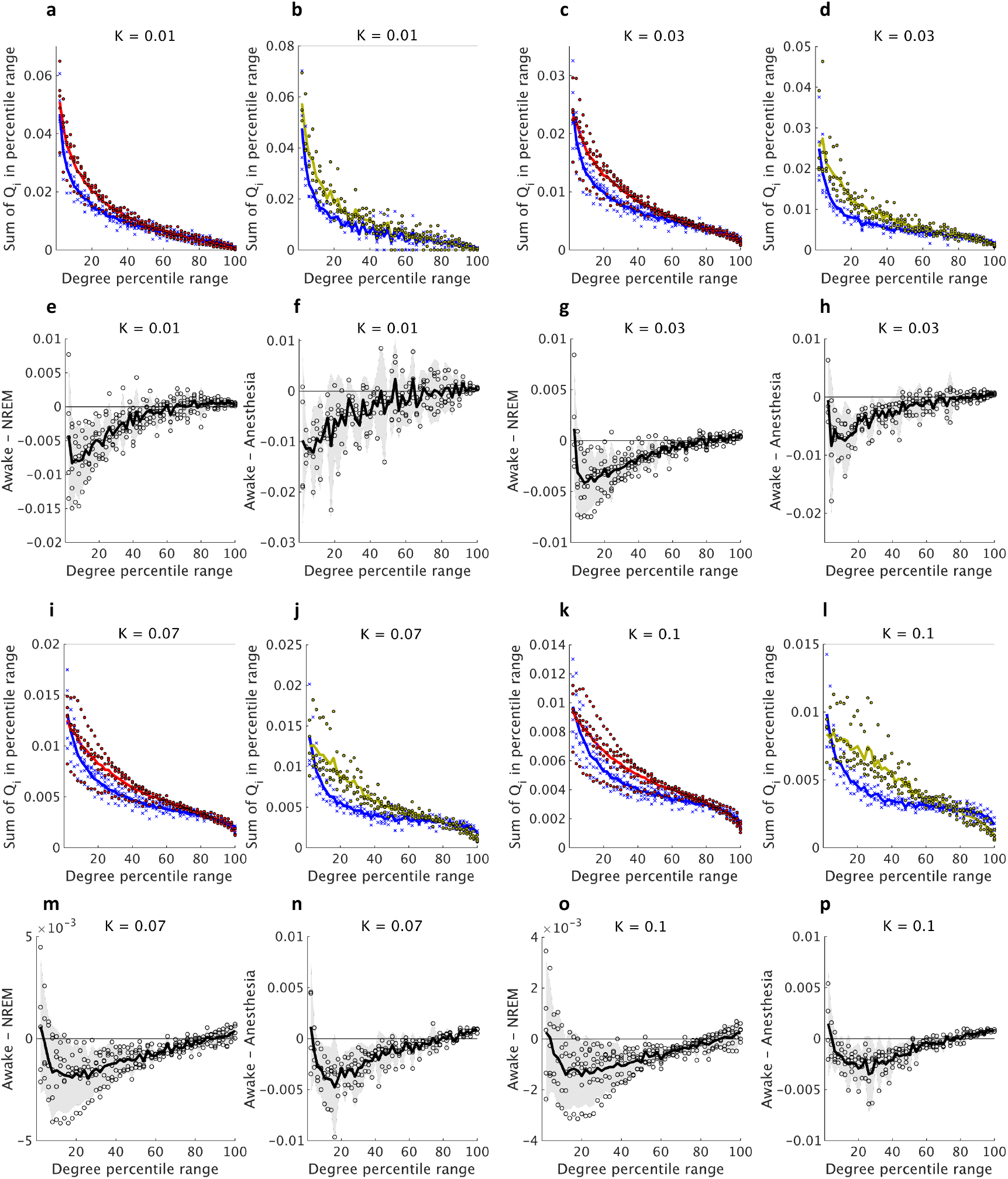
Same as Fig. 4c–f, but for different *K*.

**Figure S17:**
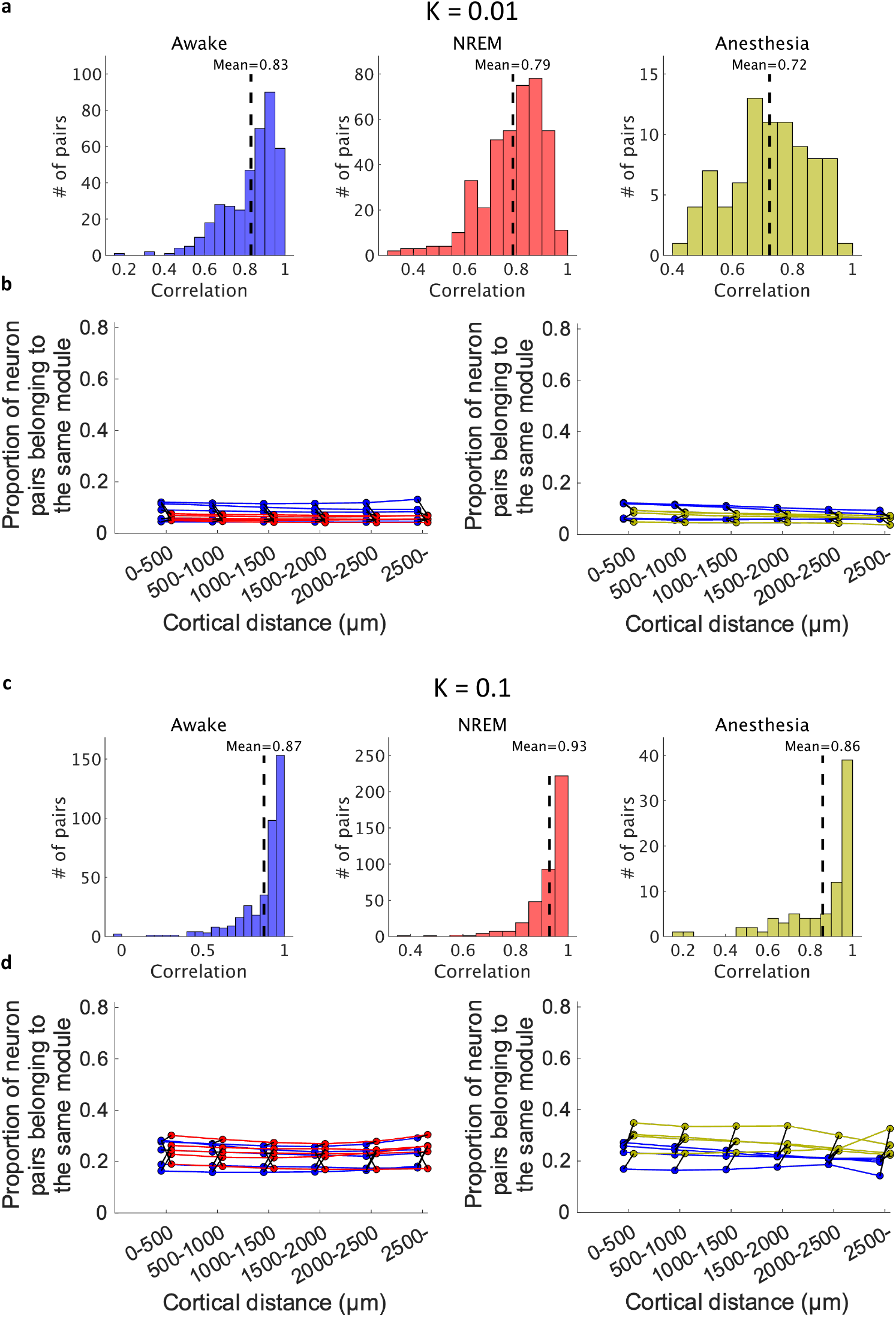
Spatially intermixed distribution of single-cell resolution functional network modules (same as Fig. 5d–h, but for different *K*).

**Figure S18:**
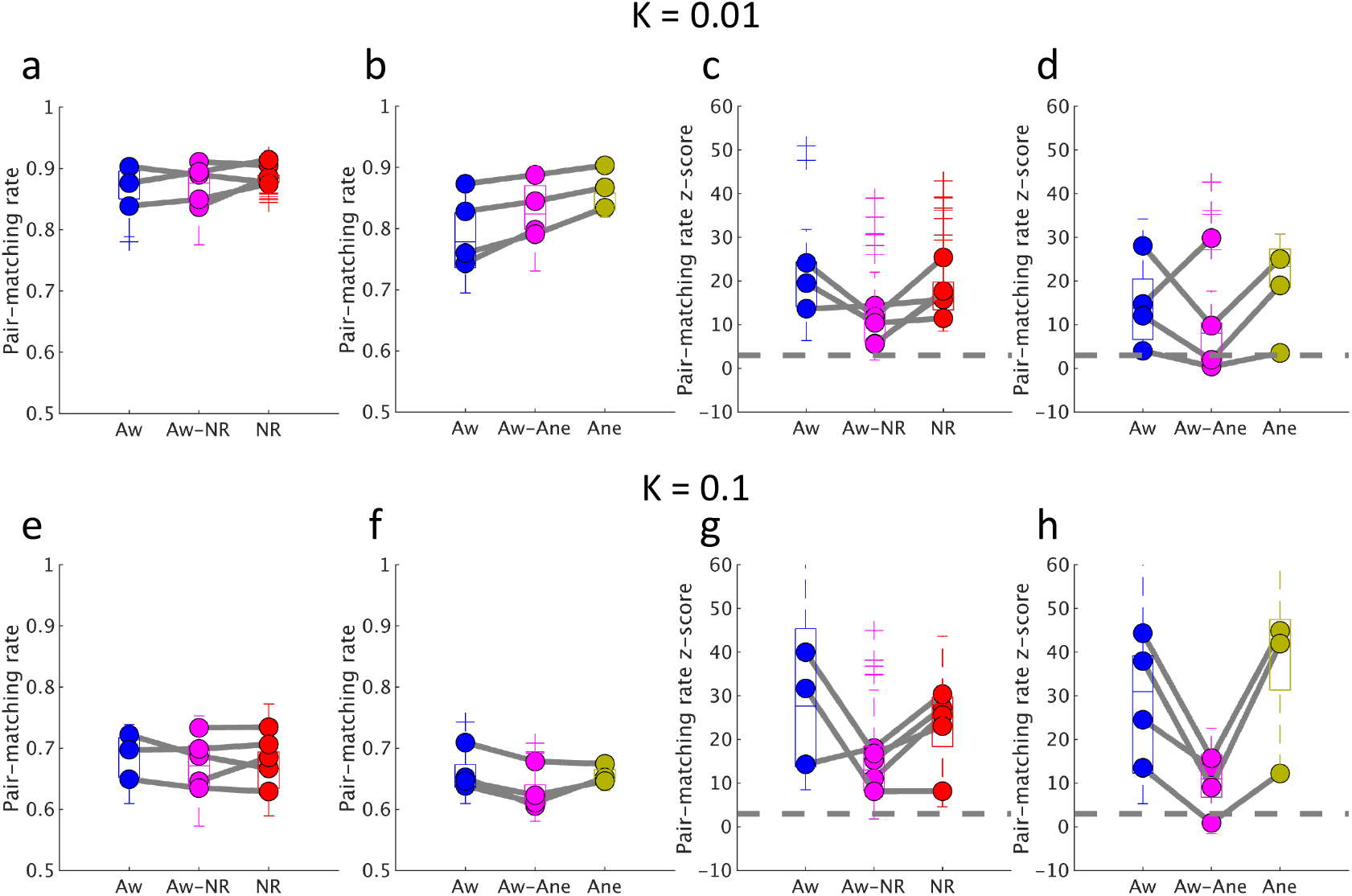
Stability of modules (same as Fig. 6, but for different *K*).

**Figure S19:**
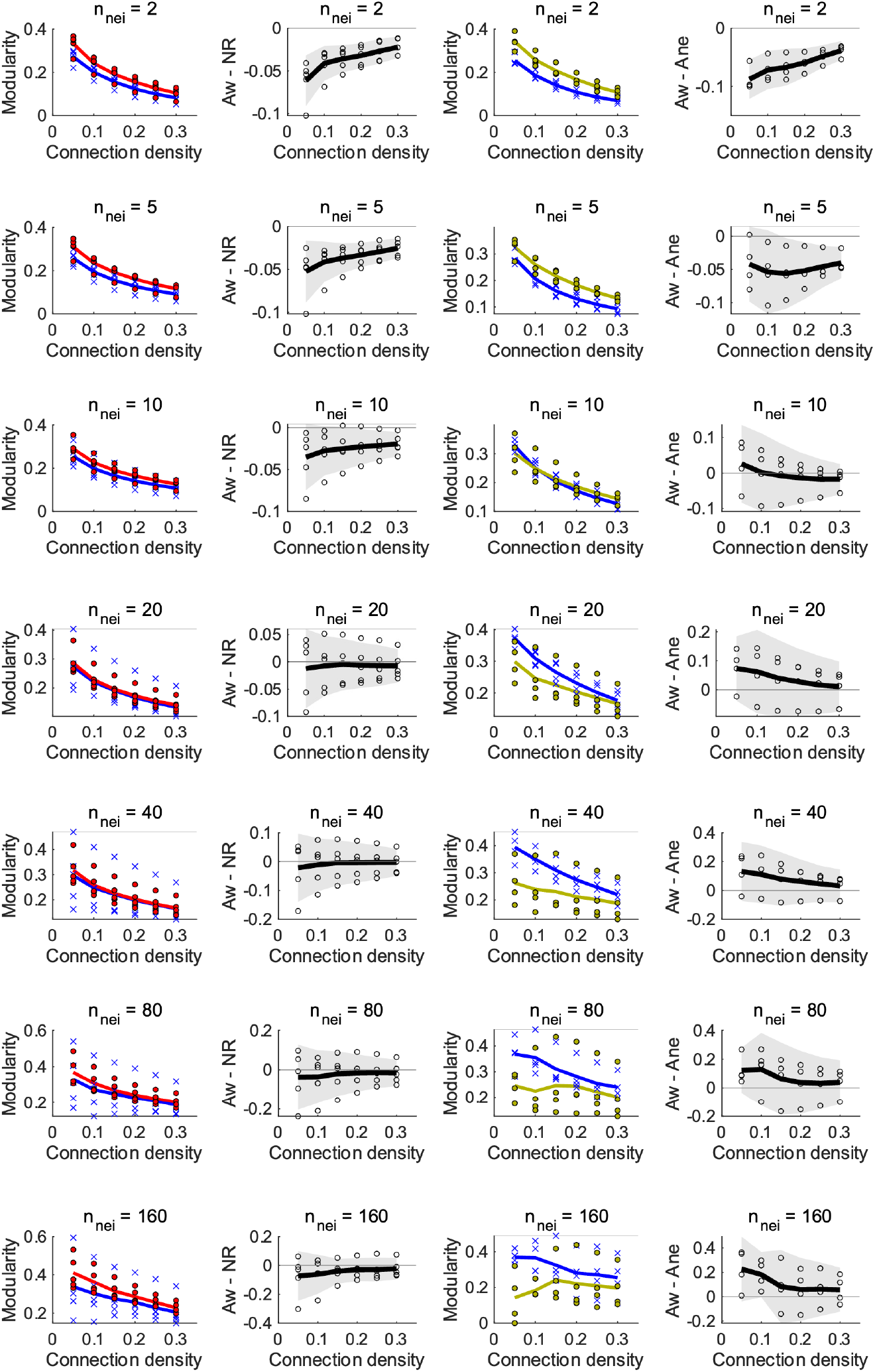
Comparison of modularity of functional networks at different spatial scales. The notation in the figure follows that of Fig. 3c–f.

**Figure S20:**
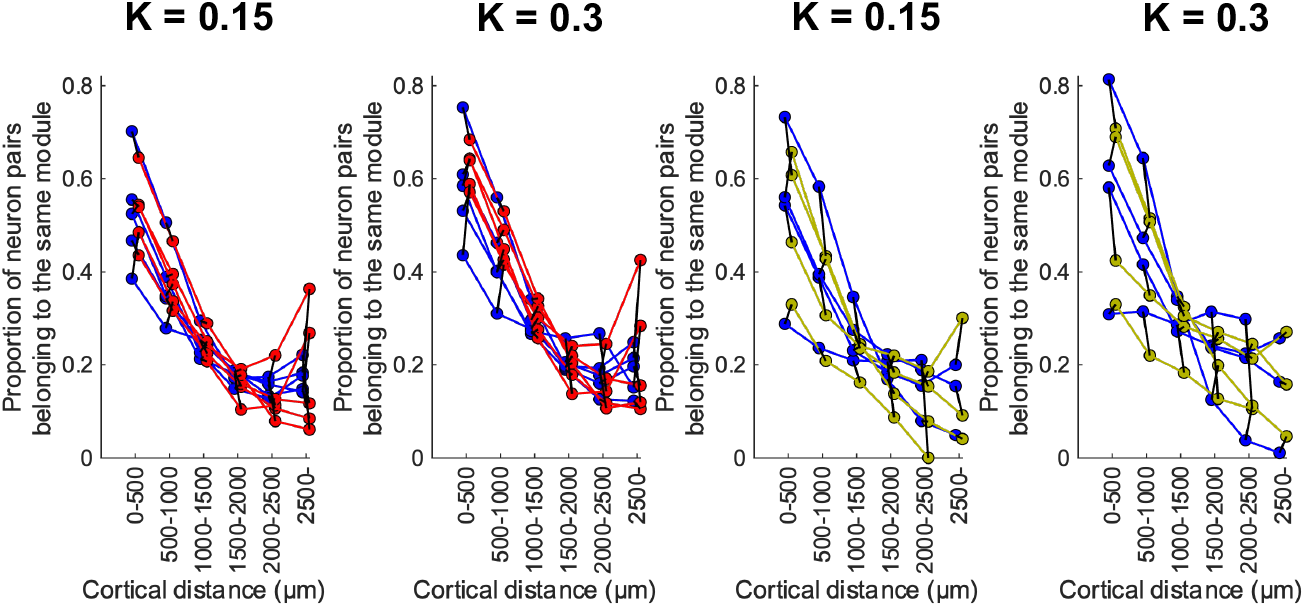
Relationships between cortical distance and module assignments of coarse-grained functional networks (same as Fig. 7g, h, but for different *K*.)

## Notes

### Competing Interest Statement

The authors have declared no competing interest.

### Summary of Updates

Abstract revised; Introduction revised; Results revised; Materials and Methods revised; Discussion revised; Supplementary Information revised; author information updated.

## References

[1] Dehaene, S. & Changeux, J.-P. Experimental and theoretical approaches to conscious processing. Neuron 70, 200–227 (2011).

[2] Mashour, G. A., Roelfsema, P., Changeux, J.-P. & Dehaene, S. Conscious processing and the global neuronal workspace hypothesis. Neuron 105, 776–798 (2020).

[3] Oizumi, M., Albantakis, L. & Tononi, G. From the phenomenology to the mechanisms of consciousness: integrated information theory 3.0. PLoS Comput. Biol. (2014).

[4] Tononi, G., Boly, M., Massimini, M. & Koch, C. Integrated information theory: from consciousness to its physical substrate. Nat. Rev. Neurosci. 17, 450–461 (2016).

[5] Deco, G., Tononi, G., Boly, M. & Kringelbach, M. L. Rethinking segregation and integration: contributions of whole-brain modelling. Nat. Rev. Neurosci. 16, 430–439 (2015).

[6] Rubinov, M. & Sporns, O. Complex network measures of brain connectivity: uses and interpretations. Neuroimage 52, 1059–1069 (2010).

[7] Seguin, C., Sporns, O. & Zalesky, A. Brain network communication: concepts, models and applications. Nat. Rev. Neurosci. (2023).

[8] Afrasiabi, M. et al. Consciousness depends on integration between parietal cortex, striatum, and thalamus. Cell Syst 12, 363–373.e11 (2021).

[9] Areshenkoff, C. N. et al. Muting, not fragmentation, of functional brain networks under general anesthesia. Neuroimage 231, 117830 (2021).

[10] Boly, M. et al. Hierarchical clustering of brain activity during human nonrapid eye movement sleep. Proc. Natl. Acad. Sci. U. S. A. 109, 5856–5861 (2012).

[11] Casali, A. G. et al. A theoretically based index of consciousness independent of sensory processing and behavior. Sci. Transl. Med. 5, 198ra105 (2013).

[12] Lee, M. et al. Connectivity differences between consciousness and unconsciousness in non-rapid eye movement sleep: a TMS-EEG study. Sci. Rep. 9, 5175 (2019).

[13] Massimini, M. et al. Breakdown of cortical effective connectivity during sleep. Science 309, 2228–2232 (2005).

[14] Schartner, M. et al. Complexity of multi-dimensional spontaneous EEG decreases during propofol induced general anaesthesia. PLoS One 10, e0133532 (2015).

[15] Schartner, M. M. et al. Global and local complexity of intracranial EEG decreases during NREM sleep. Neurosci Conscious 2017, niw022 (2017).

[16] Standage, D. et al. Dynamic reconfiguration, fragmentation, and integration of whole-brain modular structure across depths of unconsciousness. Cerebral Cortex 30, 5229–5241 (2020).

[17] Tagliazucchi, E. et al. Large-scale brain functional modularity is reflected in slow electroencephalographic rhythms across the human non-rapid eye movement sleep cycle. Neuroimage 70, 327–339 (2013).

[18] Uehara, T. et al. Efficiency of a “small-world” brain network depends on consciousness level: a resting-state FMRI study. Cereb. Cortex 24, 1529–1539 (2014).

[19] Monti, M. M. et al. Dynamic change of global and local information processing in propofol-induced loss and recovery of consciousness. PLoS Comput. Biol. 9, e1003271 (2013).

[20] Bertolero, M. A., Yeo, B. T. T., Bassett, D. S. & D’Esposito, M. A mechanistic model of connector hubs, modularity and cognition. Nat Hum Behav 2, 765–777 (2018).

[21] Bertolero, M. A., Yeo, B. T. T. & D’Esposito, M. The modular and integrative functional architecture of the human brain. Proc. Natl. Acad. Sci. U. S. A. 112, E6798–807 (2015).

[22] Chavez, M., Valencia, M., Navarro, V., Latora, V. & Martinerie, J. Functional modularity of background activities in normal and epileptic brain networks. Phys. Rev. Lett. 104, 118701 (2010).

[23] Cohen, J. R. & D’Esposito, M. The segregation and integration of distinct brain networks and their relationship to cognition. J. Neurosci. 36, 12083–12094 (2016).

[24] Gordon, E. M. et al. Three distinct sets of connector hubs integrate human brain function. Cell Rep. 24, 1687–1695.e4 (2018).

[25] Jang, H., Mashour, G. A., Hudetz, A. G. & Huang, Z. Measuring the dynamic balance of integration and segregation underlying consciousness, anesthesia, and sleep in humans. Nat Commun 15 (2024).

[26] Kim, H. et al. Estimating the integrated information measure phi from high-density electroencephalography during states of consciousness in humans. Front. Hum. Neurosci. 12, 42 (2018).

[27] Luppi, A. I. et al. Consciousness-specific dynamic interactions of brain integration and functional diversity. Nat. Commun. 10, 4616 (2019).

[28] Luppi, A. I. et al. LSD alters dynamic integration and segregation in the human brain. Neuroimage 227, 117653 (2021).

[29] Shine, J. M. et al. The dynamics of functional brain networks: Integrated network states during cognitive task performance. Neuron 92, 544–554 (2016).

[30] Tagliazucchi, E. et al. Increased global functional connectivity correlates with LSD-induced ego dissolution. Curr. Biol. 26, 1043–1050 (2016).

[31] Yanagawa, T., Chao, Z. C., Hasegawa, N. & Fujii, N. Large-scale information flow in conscious and unconscious states: an ECoG study in monkeys. PLoS One 8, e80845 (2013).

[32] Hoel, E. P., Albantakis, L. & Tononi, G. Quantifying causal emergence shows that macro can beat micro. Proc. Natl. Acad. Sci. U. S. A. 110, 19790–19795 (2013).

[33] Northoff, G. & Huang, Z. How do the brain s time and space mediate consciousness and its different dimensions? temporo-spatial theory of consciousness (TTC). Neurosci. Biobehav. Rev. 80, 630–645 (2017).

[34] Northoff, G., Wainio-Theberge, S. & Evers, K. Is temporo-spatial dynamics the “common currency” of brain and mind? in quest of “spatiotemporal neuroscience”. Phys. Life Rev. 33, 34–54 (2020).

[35] Shepherd, G. M. Foundations of the neuron doctrine (Oxford University Press, 2015).

[36] Betzel, R. F., Wood, K. C., Angeloni, C., Neimark Geffen, M. & Bassett, D. S. Stability of spontaneous, correlated activity in mouse auditory cortex. PLoS Comput. Biol. 15, e1007360 (2019).

[37] Koçillari, L. et al. Measuring stimulus-related redundant and synergistic functional connectivity with single cell resolution in auditory cortex. bioRxiv 2023.06.19.545531 (2023).

[38] Miyazaki, T. et al. Dynamics of cortical local connectivity during Sleep-Wake states and the homeostatic process. Cereb. Cortex 30, 3977–3990 (2020).

[39] Yang, W. et al. Anesthetics fragment hippocampal network activity, alter spine dynamics, and affect memory consolidation. PLoS Biol. 19, e3001146 (2021).

[40] Yu, Y., Stirman, J. N., Dorsett, C. R. & Smith, S. L. Mesoscale correlation structure with single cell resolution during visual coding. BioRxiv (2019).

[41] Clawson, W. et al. Computing hubs in the hippocampus and cortex. Sci Adv 5, eaax4843 (2019).

[42] Dann, B., Michaels, J. A., Schaffelhofer, S. & Scherberger, H. Uniting functional network topology and oscillations in the fronto-parietal single unit network of behaving primates. Elife 5 (2016).

[43] Nigam, S. et al. Rich-club organization in effective connectivity among cortical neurons. J. Neurosci. 36, 670–684 (2016).

[44] Oomoto, I. et al. Protocol for cortical-wide field-of-view two-photon imaging with quick neonatal adeno-associated virus injection. STAR Protoc 2, 101007 (2021).

[45] Ota, K. et al. Fast, cell-resolution, contiguous-wide two-photon imaging to reveal functional network architectures across multi-modal cortical areas. Neuron 109, 1810–1824.e9 (2021).

[46] Newman, M. E. J. Fast algorithm for detecting community structure in networks. Phys. Rev. E Stat. Nonlin. Soft Matter Phys. 69, 066133 (2004).

[47] Newman, M. E. J. & Girvan, M. Finding and evaluating community structure in networks. Phys. Rev. E Stat. Nonlin. Soft Matter Phys. 69, 026113 (2004).

[48] Sporns, O. & Betzel, R. F. Modular brain networks. Annu. Rev. Psychol. 67, 613–640 (2016).

[49] Bullmore, E. T. & Bassett, D. S. Brain graphs: graphical models of the human brain connectome. Annu. Rev. Clin. Psychol. 7, 113–140 (2011).

[50] Liska, A., Galbusera, A., Schwarz, A. J. & Gozzi, A. Functional connectivity hubs of the mouse brain. Neuroimage 115, 281–291 (2015).

[51] Yu, M., Sporns, O. & Saykin, A. The human connectome in alzheimer disease relationship to biomarkers and genetics. Nat. Rev. Neurol. 17, 545–563 (2021).

[52] van den Heuvel, M. P. & Sporns, O. Network hubs in the human brain. Trends Cogn. Sci. 17, 683–696 (2013).

[53] de Haan, W. et al. Disrupted modular brain dynamics reflect cognitive dysfunction in alzheimer’s disease. Neuroimage 59, 3085–3093 (2012).

[54] Yu, M. et al. Different functional connectivity and network topology in behavioral variant of frontotemporal dementia and alzheimer’s disease: an EEG study. Neurobiol. Aging 42, 150–162 (2016).

[55] Buckner, R. L. et al. Cortical hubs revealed by intrinsic functional connectivity: mapping, assessment of stability, and relation to alzheimer’s disease. J. Neurosci. 29, 1860–1873 (2009).

[56] Crossley, N. A. et al. The hubs of the human connectome are generally implicated in the anatomy of brain disorders. Brain 137, 2382–2395 (2014).

[57] Filippi, M. et al. Changes in functional and structural brain connectome along the alzheimer’s disease continuum. Mol. Psychiatry 25, 230–239 (2020).

[58] Yu, M. et al. Selective impairment of hippocampus and posterior hub areas in alzheimer’s disease: an MEG-based multiplex network study. Brain 140, 1466–1485 (2017).

[59] Ohki, K., Chung, S., Ch’ng, Y. H., Kara, P. & Reid, R. C. Functional imaging with cellular resolution reveals precise micro-architecture in visual cortex. Nature 433, 597–603 (2005).

[60] Peron, S. P., Freeman, J., Iyer, V., Guo, C. & Svoboda, K. A cellular resolution map of barrel cortex activity during tactile behavior. Neuron 86, 783–799 (2015).

[61] Muldoon, S. F., Soltesz, I. & Cossart, R. Spatially clustered neuronal assemblies comprise the microstructure of synchrony in chronically epileptic networks. Proceedings of the National Academy of Sciences 110, 3567–3572 (2013).

[62] Han, Y. et al. The logic of single-cell projections from visual cortex. Nature 556, 51–56 (2018).

[63] O’Rawe, J. F. et al. Excitation creates a distributed pattern of cortical suppression due to varied recurrent input. Neuron (2023).

[64] Sanzeni, A. et al. Mechanisms underlying reshuffling of visual responses by optogenetic stimulation in mice and monkeys. Neuron 111, 4102–4115.e9 (2023).

[65] Fişek, M. et al. Cortico-cortical feedback engages active dendrites in visual cortex. Nature (2023).

[66] Izquierdo-Serra, M., Hirtz, J. J., Shababo, B. & Yuste, R. Two-photon optogenetic mapping of excitatory synaptic connectivity and strength. iScience 8, 15–28 (2018).

[67] Betzel, R. F. & Bassett, D. S. Multi-scale brain networks. Neuroimage 160, 73–83 (2017).

[68] Betzel, R. F. Organizing principles of whole-brain functional connectivity in zebrafish larvae. Netw Neurosci 4, 234–256 (2020).

[69] Leung, A. & Tsuchiya, N. Emergence of integrated information at macro timescales in real neural recordings. Entropy 24, 625 (2022).

[70] Meshulam, L., Gauthier, J. L., Brody, C. D., Tank, D. W. & Bialek, W. Coarse graining, fixed points, and scaling in a large population of neurons. Phys. Rev. Lett. 123, 178103 (2019).

[71] Munn, B. R. et al. Multiscale organization of neuronal activity unifies scale-dependent theories of brain function. Cell 0 (2024).

[72] Rubinov, M., Ypma, R. J. F., Watson, C. & Bullmore, E. T. Wiring cost and topological participation of the mouse brain connectome. Proc. Natl. Acad. Sci. U. S. A. 112, 10032–10037 (2015).

[73] Rosas, F. E. et al. Reconciling emergences: An information-theoretic approach to identify causal emergence in multivariate data. PLoS Comput. Biol. 16, e1008289 (2020).

[74] Tononi, G., Sporns, O. & Edelman, G. M. A measure for brain complexity: relating functional segregation and integration in the nervous system. Proc. Natl. Acad. Sci. U. S. A. 91, 5033–5037 (1994).

[75] Varley, T. F. & Hoel, E. Emergence as the conversion of information: a unifying theory. Philos. Trans. A Math. Phys. Eng. Sci. 380, 20210150 (2022).

[76] Herbet, G. & Duffau, H. Revisiting the functional anatomy of the human brain: Toward a meta-networking theory of cerebral functions. Physiol. Rev. 100, 1181–1228 (2020).

[77] Posner, M. I., Petersen, S. E., Fox, P. T. & Raichle, M. E. Localization of cognitive operations in the human brain. Science 240, 1627–1631 (1988).

[78] Manita, S. et al. A top-down cortical circuit for accurate sensory perception. Neuron 86, 1304–1316 (2015).

[79] Miyamoto, D. et al. Top-down cortical input during NREM sleep consolidates perceptual memory. Science 352, 1315–1318 (2016).

[80] Saito, Y. et al. Amygdalo-cortical dialogue underlies memory enhancement by emotional association. Neuron 0 (2025).

[81] Kaplan, H. S. & Zimmer, M. Brain-wide representations of ongoing behavior: a universal principle? Curr. Opin. Neurobiol. 64, 60–69 (2020).

[82] Musall, S., Kaufman, M. T., Juavinett, A. L., Gluf, S. & Churchland, A. K. Single-trial neural dynamics are dominated by richly varied movements. Nat. Neurosci. 22, 1677–1686 (2019).

[83] Salkoff, D. B., Zagha, E., McCarthy, E. & McCormick, D. A. Movement and performance explain widespread cortical activity in a visual detection task. Cereb. Cortex 30, 421–437 (2020).

[84] Stringer, C. et al. Spontaneous behaviors drive multidimensional, brainwide activity. Science 364, 255 (2019).

[85] Aimon, S. et al. Fast near-whole -brain imaging in adult drosophila during responses to stimuli and behavior. PLoS Biol. 17, e2006732 (2019).

[86] Kato, S. et al. Global brain dynamics embed the motor command sequence of caenorhabditis elegans. Cell 163, 656–669 (2015).

[87] Westlin, C. et al. Improving the study of brain-behavior relationships by revisiting basic assumptions. Trends Cogn. Sci. (2023).

[88] Shiba, Y. et al. Allogeneic transplantation of iPS cell-derived cardiomyocytes regenerates primate hearts. Nature 538, 388–391 (2016).

[89] Kobayashi, K. et al. Survival of corticostriatal neurons by rho/rho-kinase signaling pathway. Neurosci. Lett. 630, 45–52 (2016).

[90] Kawai, S., Takagi, Y., Kaneko, S. & Kurosawa, T. Effect of three types of mixed anesthetic agents alternate to ketamine in mice. Exp. Anim. 60, 481–487 (2011).

[91] Tsunematsu, T. et al. Acute optogenetic silencing of orexin/hypocretin neurons induces slow-wave sleep in mice. J. Neurosci. 31, 10529–10539 (2011).

[92] Tsunematsu, T. et al. Long-lasting silencing of orexin/hypocretin neurons using archaerhodopsin induces slow-wave sleep in mice. Behav. Brain Res. 255, 64–74 (2013).

[93] Saito, Y. et al. Emotional association enhances perceptual memory through amygdalo-cortical inputs during NREM sleep. bioRxiv 2023.05.23.541852 (2023).

[94] Pnevmatikakis, E. A. & Giovannucci, A. NoRMCorre: An online algorithm for piecewise rigid motion correction of calcium imaging data. J. Neurosci. Methods 291, 83–94 (2017).

[95] Ito, T. et al. Low computational-cost cell detection method for calcium imaging data. Neurosci. Res. 179, 39–50 (2022).

[96] Dombeck, D. A., Khabbaz, A. N., Collman, F., Adelman, T. L. & Tank, D. W. Imaging large-scale neural activity with cellular resolution in awake, mobile mice. Neuron 56, 43–57 (2007).

[97] Pachitariu, M. et al. Suite2p: beyond 10,000 neurons with standard two-photon microscopy. bioRxiv 061507 (2016).

[98] Friedrich, J., Zhou, P. & Paninski, L. Fast online deconvolution of calcium imaging data. PLoS Comput. Biol. 13, e1005423 (2017).

[99] Schneider, C. A., Rasband, W. S. & Eliceiri, K. W. NIH image to ImageJ: 25 years of image analysis. Nat. Methods 9, 671–675 (2012).

[100] Newman, M. E. J. Modularity and community structure in networks. Proc. Natl. Acad. Sci. U. S. A. 103, 8577–8582 (2006).

[101] Blondel, V. D., Guillaume, J. L., Lambiotte, R. & others. Fast unfolding of communities in large networks. Journal of statistical (2008).

[102] Guimer’a, R. & Amaral, L. A. N. Cartography of complex networks: modules and universal roles. J. Stat. Mech. 2005, nihpa35573 (2005).

[103] Guimer’a, R. & Nunes Amaral, L. A. Functional cartography of complex metabolic networks. Nature 433, 895–900 (2005).

[104] Latora, V. & Marchiori, M. Efficient behavior of small-world networks. Phys. Rev. Lett. 87, 198701 (2001).

[105] Newman, M. E. J. The structure and function of complex networks. SIAM Rev. 45, 167–256 (2003).

